# LysM-RLK plays an ancestral symbiotic function in plants

**DOI:** 10.1101/2024.01.16.575821

**Authors:** Eve Teyssier, Sabine Grat, David Landry, Mathilde Ouradou, Melanie K. Rich, Sebastien Fort, Jean Keller, Benoit Lefebvre, Pierre-Marc Delaux, Malick Mbengue

## Abstract

Arbuscular mycorrhiza (AM) with soilborne Glomeromycota fungi was pivotal in the conquest of land by plants almost half a billion years ago. In flowering plants, it is hypothesised that AM is initiated by the perception of AM-fungi-derived chito- and lipochito-oligosaccharides (COs/LCOs) in the host via Lysin Motif Receptor-Like Kinases (LysM-RLKs). However, it remains uncertain whether plant perception of these molecules is a prerequisite for AM establishment and for its origin. Here, we made use of the reduced LysM-RLK complement present in the liverwort *Marchantia paleacea* to assess the conservation of the role played by this class of receptors during AM and in COs/LCOs perception. Our reverse genetic approach demonstrates the critical function of a single LysM-RLK, LYKa, in AM formation, thereby supporting an ancestral function for this receptor in symbiosis. Binding studies, cytosolic calcium variation recordings and genome-wide transcriptomics indicate that another LysM-RLK of *M. paleacea*, LYR, is also required for triggering a response to COs/LCOs, despite being dispensable for AM formation. Collectively, our results demonstrate that the perception of symbionts by LysM-RLK is an ancestral feature in land plants, and suggest the existence of yet-uncharacterised AM-fungi signals.

## Introduction

To ensure their water and mineral nutrition, most land plants form arbuscular mycorrhiza (AM) with soil-borne Glomeromycota fungi [1]. This ∼450 million years old symbiosis was key in driving land colonisation by plants [2]. In angiosperms, AM is thought to be initiated via the perception by the host plant of AM-fungi derived chito-(COs) and lipochito-oligosaccharides (LCOs) leading to the activation of a conserved signalling pathway referred to as the Common Symbiosis Signalling Pathway (CSSP) [3]. Mutation of any CSSP gene, such as the receptor-like kinase *SYMRK* [4], leads to the inability to associate with AM-fungi in species as diverse as legumes [4,5], monocots [6,7] or bryophytes [8,9].

Genetics in legumes and monocots have demonstrated that members of the diversified Lysin motif Receptor-Like Kinase (LysM-RLK) family, which encompasses up to 22 members in legumes [10], are important for the perception of these AM-fungi derived molecules [11,12]. Only one double mutant in the legumes *Medicago truncatula* and *Lotus japonicus* showed a phenotype phenocopying mutants of the CSSP with a total lack of colonisation, demonstrating the importance of this family of receptors for AM in flowering plants [13,14]. However, none of the mutant combinations fully abolished the perception of both COs and LCOs, thus not allowing to determine whether these symbiotic signals are sufficient for AM establishment or whether other signal(s) are involved. The LysM-RLK family expansion that occurred in angiosperms likely complicate such analysis due to genetic redundancy.

Angiosperms, such as legumes and monocots, belong to the vascular plants. The other group of land plants, the bryophytes, includes species with a haploid-dominant life-cycle such as the liverwort *Marchantia paleacea*. Because the bryophyte and vascular plant lineages diverged from each other soon after the colonisation of lands by plants, any trait found to be conserved in both clades can be inferred as ancestral in plants [15,16]. Using reverse genetics in *M. paleacea* and comparison with vascular plants, it has been demonstrated that AM has been relying on lipids provided by the host plants for 450 million years [17]. Furthermore, mutations in any of the CSSP gene in *M. paleacea* lead to the total absence of colonization by AM fungi, as observed in angiosperms [8,9]. It can thus be hypothesised that at least part of the signalling processes involved in the formation of AM in *M. paleacea* and angiosperms are similar. In contrast to angiosperms, the liverwort *M. paleacea* contains only four LysM-RLKs (see below), limiting the expected genetic redundancy observed in angiosperms. *M. paleacea* therefore represents a unique model to study the role of CO- and LCOs-signalling and determine whether other signals are involved in AM establishment.

In this study, we demonstrate the essential role for AM in *M. paleacea* of a single LysM-RLK, pro- ortholog to the highly diversified CERK1 clade in most angiosperms. Moreover, we demonstrate that the ability of *M. paleacea* to respond to COs and LCOs is fully abolished in that mutant. Yet, we present genetic evidence that signalling triggered by COs and LCOs is not essential for AM in *M. paleacea*, suggesting the existence of additional and uncharacterised AM-fungi signals.

## Results

### *Marchantia paleacea* has a limited number of LysM-RLKs

Phylogenetic analyses indicate that LysM-RLK originated in streptophyte green algae and diversified following the colonisation of land by plants [18,19]. Extant liverworts harbour four clades, including three LYKs with a predicted active kinase and one LYR with a deleted activation loop in the kinase domain [20]. All LysM-RLKs of *M. paleacea* contain a predicted N-terminal signal peptide, a LysM-containing extracellular domain, a transmembrane region and an intracellular kinase domain. Based on the established nomenclature for LysM-RLKs [10], *M. paleacea* LYKa belongs to the LYK-I subclade that contains the immunity-related chitin co-receptor CERK1 from *Arabidopsis thaliana* [21] and the symbiotic Nod factor co-receptor NFR1 from *Lotus japonicus* [22] (**Fig 1**). As a unique member, MpaLYKa therefore represents the pro-ortholog in *M. paleacea* of all members of the LYK-I subclade in angiosperm. Similarly, MpaLYKb and MpaLYKc are pro-orthologous to the LYK-II and LYK-III subclades, respectively, while the single MpaLYR is pro-orthologous to the entire LYR-I to LYR-IV subclades (**Fig 1**). *Marchantia polymorpha*, that has lost the ability to form AM [23], contains only two *LYKs* and one *LYR* [20]. Remnants of a third *LYK* gene in *M. polymorpha*, encoding a truncated receptor from the LYK-II subclade, accounts for this discrepancy between Marchantia species (**Fig 1**).

**Fig 1.**
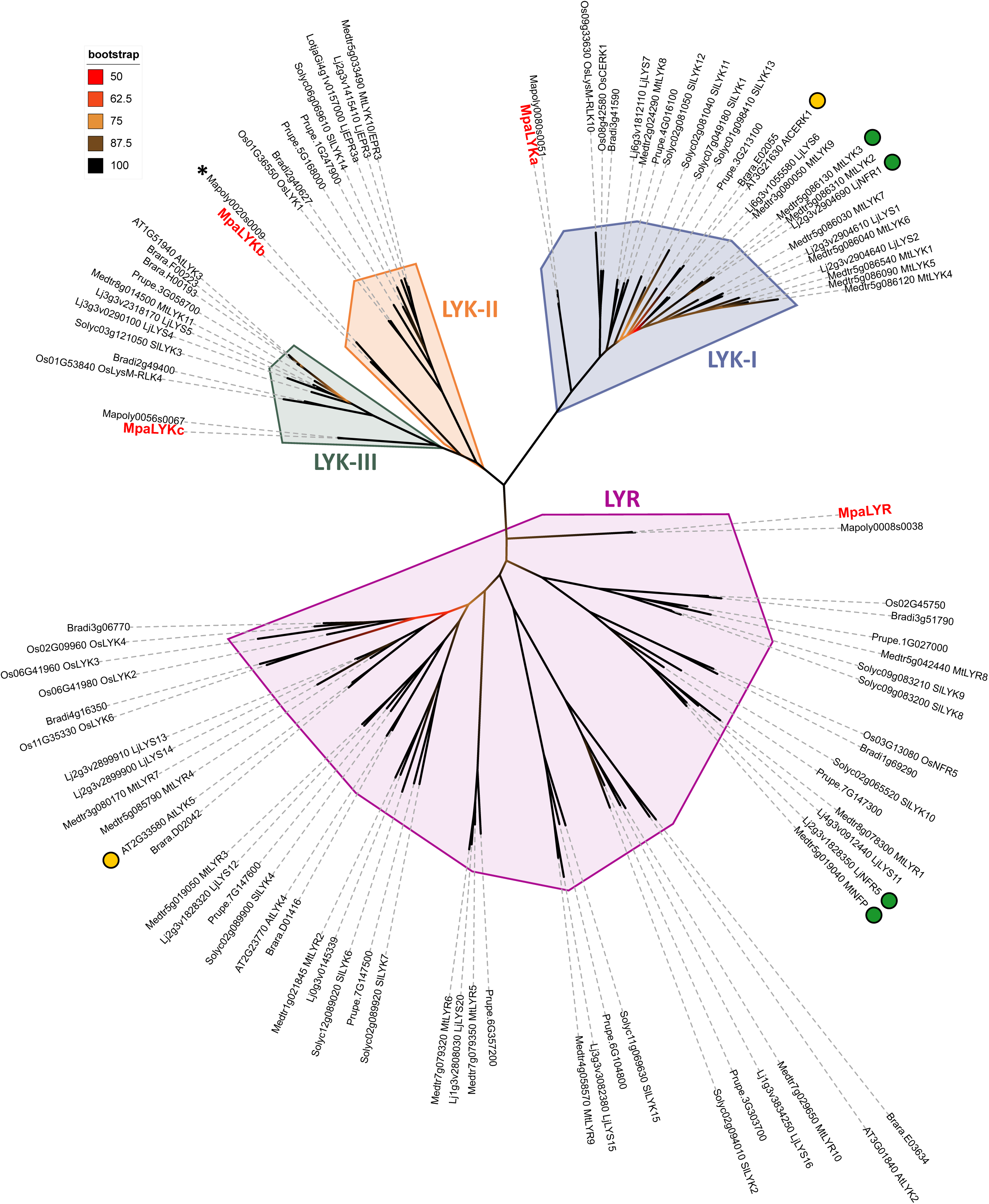
*Marchantia paleacea* possesses four LysM Receptor-Like Kinases. Phylogenetic analysis of 102 LysM-RLKs from land plant species. Full-length protein sequences from *A. thaliana* (AT)*, S. lycopersicum* (Solyc), *O. sativa* (Os), *P. persica* (Prupe), *L. japonicus* (Lj), *M. truncatula* (Medtr), *M. polymorpha* (Mapoly) and *M. paleacea* (Marpal) were aligned with MUSCLE prior unrooted Maximum Likelihood phylogenetic tree construction using 1000 bootstraps resampling value. Branch colours indicate bootstrap values. Four phylogenetic subclades are highlighted: LYK-I (blue), LYK-II (orange), LYK-III (green), LYR (pink). *M. paleacea* proteins are labelled in red. The star sign indicates the truncated LYKb homolog in *M. polymorpha*. Notable receptors for long-chain CO perception in Arabidopsis or Nod factors perception in legumes are indicated with yellow or green dots, respectively.

### Mutation of a single *LYK* gene in *Marchantia paleacea* leads to symbiosis impairment

To determine the contribution of individual LysM-RLKs from the *M. paleacea* LYK clade to AM, we generated loss-of-function mutants for *MpaLYKa*, *MpaLYKb* or *MpaLYKc*. For that, CRISPR/Cas9 was used in combination with single guide-RNAs (sgRNA) designed to target the 5’ region of the genes, hereby breaking the link between the signal peptide and the remaining part of these type-I membrane proteins (**S1 Fig**). Following plant transformation and subsequent progeny selection, we recovered independent mutant lines for each *LYK* gene. Sanger sequencing of induced mutations are summarized in **S1 Table**. For *MpaLYKa* and *MpaLYKb*, only one of the two sgRNAs proved effective in inducing frameshift-causing INDELs whereas the two different sgRNAs designed against *MpaLYKc* were effective in inducing *null* mutations. For each LYK, we kept a minimum of two *null* mutant lines resulting from independent T-DNA transformation events and assessed their ability to form AM in comparison to an empty vector-containing control line. Six weeks after inoculation with spores of the AM fungus *Rhizophagus irregularis*, we evaluated the colonisation status of *M. paleacea* by histological observations of transversal (**Fig 2A-D**) or longitudinal (**S2 Fig A-D**) sections of the thallus. A strong pigmentation along the midrib of the thallus is a marker of a positive mycorrhizal colonisation in *M. paleacea* [24]. This pigmentation was visible in control, *lykb* and *lykc* mutants but absent in *lyka*, suggesting successful symbiosis establishment in *lykb* and *lykc* and a lack of fungal colonisation in *lyka* (**Figs 2A-D and S2 A-D - middle panels**). Wheat-Germ-Agglutinin (WGA)-Alexa Fluor staining of fungal cell wall confirmed the presence of arbuscules in the thalli of control, *lykb* and *lykc* plants. In contrast, no intercellular hyphae nor arbuscules were observed in *lyka* (**Figs 2A-D and S2A-D - left and right panels**), confirming the absence of AM in these mutants. Over two large scale experiments, quantitative analysis of the exclusion zone length [2] as well as the percentages of mycorrhizal plants per genotype did not reveal differences between *lykb* or *lykc* alleles compared to the control (**Figs 2E and S2E**). In contrast, these experiments confirmed AM-defectiveness in three independent *lyka* mutants, representing approximately 300 measurements in total for this class of mutant.

**Fig 2.**
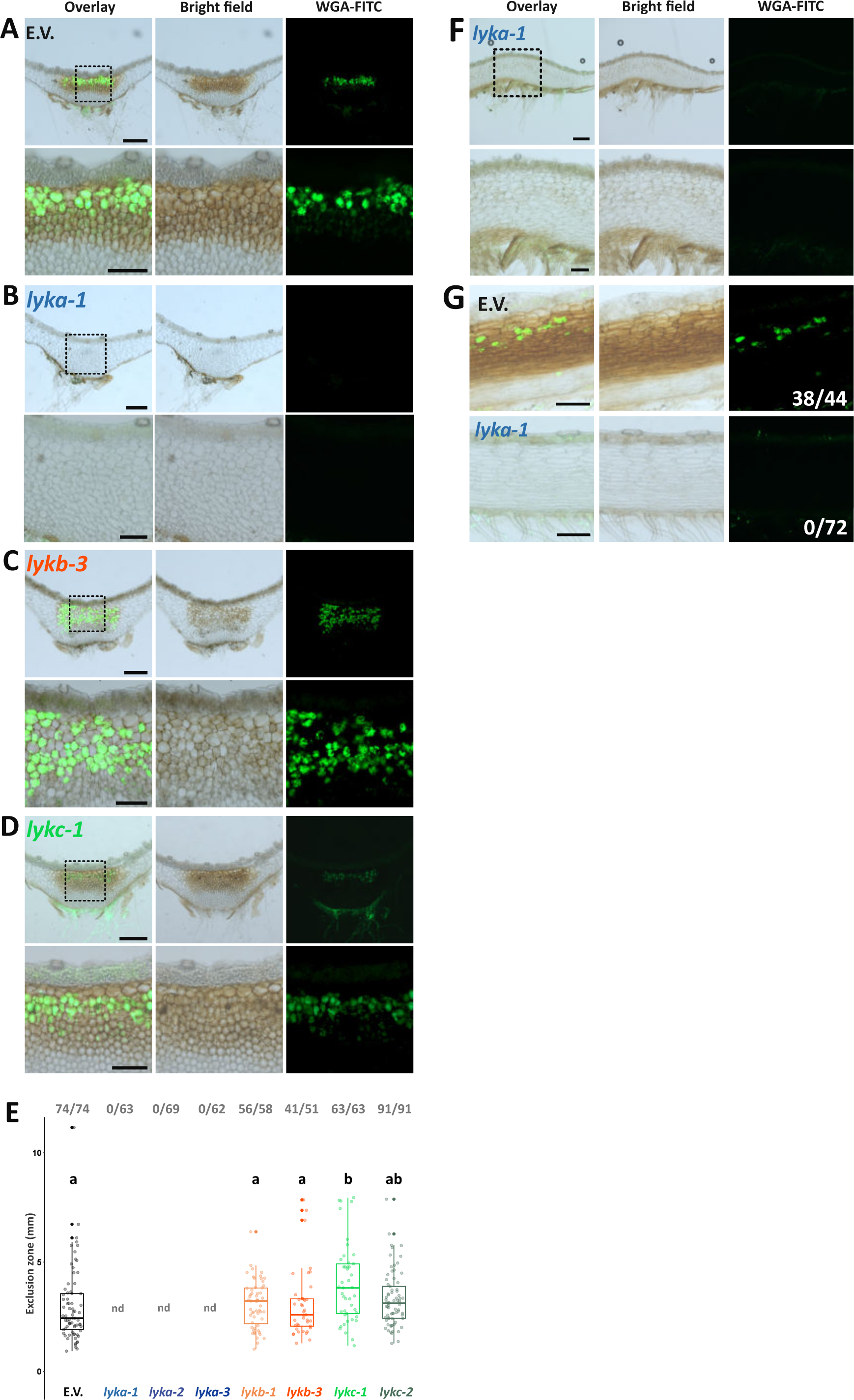
*MpaLYKa* is essential for arbuscular mycorrhiza in *M. paleacea*. (**A - D**) Representative images of transversal sections of *M. paleacea* control (E.V.) or *lyk* mutants, as indicated, six weeks after inoculation with *Rhizophagus irregularis*. Left panels are overlays of bright fields (middle panels) and fluorescent (right panels) images. Wheat germ agglutinin (WGA) coupled to Alexa Fluor 488 was used to detect fungal structures. Dashed line delimited insets are enlarged underneath the original images. Scale bars are 500 µm and 200 µm for insets. (**E**) Quantitative analysis of the exclusion zone length on mycorrhizal plants for control (E.V.) and two independent loss-of-function mutants for each *LYK*. Fractions in bold grey represent mycorrhizal thalli over total thalli assessed for each genotype. Different letters indicate differences to control inferred by ANOVA followed by Tukey’s HSD post-hoc test (p-value < 0.05). Control (E.V.) values are shared with Fig 3 and statistical analysis was performed on the whole dataset. “nd” stands for not determined. (**F**) Longitudinal sections of the *lyka-1* mutant 10 weeks after inoculation with *Rhizophagus irregularis*. (**G**) Longitudinal sections of *M. paleacea* control (E.V.) or a *lyka-1* mutant six weeks after co-cultivation in close proximity with wild-type mycorrhized *M. paleacea* nurse plants. Numbers of mycorrhized thalli over total thalli tested are indicated. Scale bar = 200 µm.

To verify that the absence of fungal colonisation in *lyka* was not due to a delay in AM establishment, we harvested *lyka* ten weeks after inoculation with *R. irregularis* spores. Again, we were unable to observe accumulation of the AM-related pigment or positive WGA-Alexa Fluor signal in the thalli of inoculated *lyka* plants (**Fig 2F**). Next, we tested for complementation of *lyka* when co-cultivated in proximity to wild-type *M. paleacea* that had been hosting AM fungi for several weeks. This nurse-plant method has been successfully used to distinguish between defaults in penetration and in survival of AM fungi in mutants. Six weeks after the start of the co-culture with nurse plants, the empty-vector control plants showed an 86% success rate in fungal colonisation (**Fig 2G**). Conversely, none of the *lyka* plants grown under these conditions were colonised (**Fig 2G**), indicating that the *lyka* phenotype is not due to a secondary effect on the fungal symbiont but is rather the direct consequence of a non-functional MpaLYKa-dependent signalling pathway in the host. We therefore conclude that, within the *LYK* clade of *M. paleacea*, *MpaLYKa* is critical for AM establishment whereas *MpaLYKb* and *MpaLYKc* are dispensable.

### The single LYR gene from *Marchantia paleacea* is not essential for arbuscular mycorrhiza

Contrary to AM-host angiosperms, which carry between 4 and 11 LYR paralogs [10], the genome of *M. paleacea* contains a single *LYR* gene [23] (**Fig 1**). To assess its contribution to symbiosis, we generated *lyr* loss-of-function alleles using CRISPR/Cas9 and two independent sgRNAs, as previously described (**Fig S1 and Table S1**), and tested their ability to form AM. As performed for the *lyk* mutants, we evaluated the colonisation status of the *lyr* mutants by histological observations of transversal (**Fig 3A**) or longitudinal (**Fig S3A**) sections of the thallus. We observed no differences in thallus pigmentation between empty-vector control and *lyr*, suggesting similar colonisation levels in both genotypes. WGA-staining confirmed the presence of arbuscules in both control and *lyr* lines (**Figs 3A and S3A**). Similarly to what we observed for the *lykb* and *lykc* mutants, quantitative analysis of the exclusion zone length [2] as well as the percentages of mycorrhizal plants per genotype did not point at reproducible differences between *lyr* alleles and control (**Figs 3B and S3B**). Over two large scale experiments, control plants displayed a total of 98% mycorrhizal success rate and *lyr* alleles followed closely with 92% and 96% mycorrhizal success rate. The mean lengths of the exclusion zone were similar between genotypes, although a slight difference of approx. 1 mm was measured across replicates (**Figs 3B and S3B**). The absence of AM phenotype in mutants of the single *LYR* gene demonstrates the non-essential role for this clade of LysM-RLK for AM establishment in *M. paleacea*.

**Fig 3.**
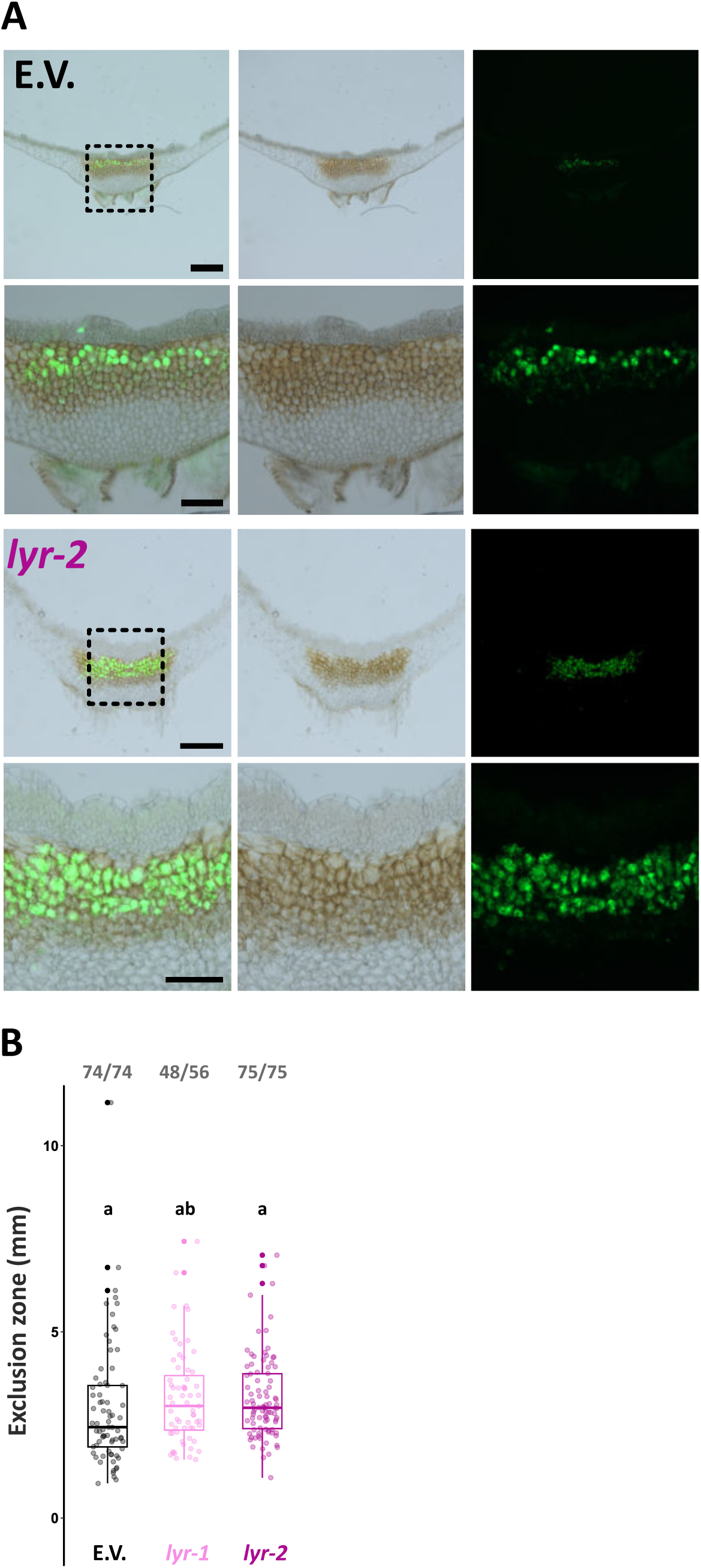
*MpaLYR* is dispensable for arbuscular mycorrhiza in *M. paleacea*. **(A)** Representative images of transversal sections of *M. paleacea* control (E.V.) or the *lyr-2* mutant six weeks after inoculation with *Rhizophagus irregularis*. Left panels are overlays of bright field (middle panels) and fluorescent (right panels) images. Wheat germ agglutinin (WGA) coupled to Alexa Fluor 488 was used to detect fungal structures. Dashed line delimited insets are enlarged underneath the original images. Scale bars are 500 µm and 200 µm for insets. (**B**) Quantitative analysis of the exclusion zone length on mycorrhizal plants for control (E.V.) and two *lyr* mutants. For each genotype, fractions in bold grey represent mycorrhizal thalli over total thalli assessed. Different letters indicate differences to control inferred by ANOVA followed by Tukey’s HSD post-hoc test (p-value < 0.05). Control values are shared with Fig 2 and statistical analysis was performed on the whole dataset.

### MpaLYR is the COs/LCOs receptor of *Marchantia paleacea*

In order to gain insights into the functional roles of *M. paleacea* LysM-RLKs in the perception of COs and LCOs, we conducted binding assays using tagged receptors transiently expressed in *Nicotiana benthamiana* cells. Firstly, the sub-cellular localization of the four receptor proteins tagged in their C-terminus with mCherry was verified by confocal microscopy when co-expressed with the well-known plasma-membrane resident Arabidopsis Flagellin Sensing 2 (FLS2) receptor which was fused with GFP [25,26] (**Fig 4A**). For each receptor, the mCherry signal was visible on the epidermal cell periphery and perfectly superimposed with the GFP signal, demonstrating the correct trafficking for all four *M. paleacea* LysM-RLK fusion constructs to the plasma membrane in *N. benthamiana* (**Fig 4A**). Next, microsomal preparations from leaf tissue expressing single LysM-RLK were mixed with increasing concentrations of cross-linkable biotinylated chito-pentaose or chito-heptaose (hereafter CO5* and CO7*, respectively) before detection of the biotin moieties and the mCherry tag by Western blot (**Fig 4B and D**). While none of the LYK receptors exhibited binding to CO5* or CO7* at concentrations up to 1 μM, a clear signal was observed for MpaLYR when challenged with only 100 nM of CO5* or CO7*. (**Fig 4B and D**). The over-exposure of the streptavidin blots did not reveal any weak signals, thereby suggesting that neither MpaLYKa, MpaLYKb nor MpaLYKc are able to bind CO5* or CO7* in these conditions (**Fig S4**). To determine the affinity of the MpaLYR receptor with regards to COs, experiments were conducted in which various concentrations of CO5* and CO7*, ranging from 1 nM to 2 μM, were tested (**Fig 4C and E**). The presence of ligand-bound LYR proteins was observed at concentrations as low as 10 nM of CO5* or 50 nM of CO7* (**Fig 4C and E**). Saturation of the signal between 250 and 500 nM indicate that the LYR receptor exhibits a high affinity for both short and long CO molecules. To confirm these findings and test whether MpaLYR could also bind LCOs, a competition assay was performed wherein MpaLYR-containing microsomes were simultaneously incubated with 100 nM CO5* and 10-fold higher concentration (1 µM) of unlabelled chito-tetraose (CO4), CO7, or the major forms of lipo-chitooligosaccharides produced by *R. irregularis*, LCO-V(C18:1, Fuc/MeFuc), hereafter Fuc/MeFuc-LCO [27] (**Fig 4F**). Additionally, 1 µM of de-acetylated chitin (chitosan) fragments or 10 ng.µL^-1^ peptidoglycan fragments derived from *Bacillus subtilis* were used as negative controls. The application of unlabelled CO4, CO7 or Fuc/MeFuc-LCOs resulted in a strong reduction in the binding of CO5* to MpaLYR when compared to control condition in the absence of competitors. In contrast, the application of bacterial peptidoglycan or chitosan fragments had no discernible effect on the binding of CO5* to the MpaLYR receptor (**Fig 4F**). Collectively, these results support the conclusion that, within the LysM-RLK family of *M. paleacea*, MpaLYR serves as the CO and LCO receptor.

**Fig 4.**
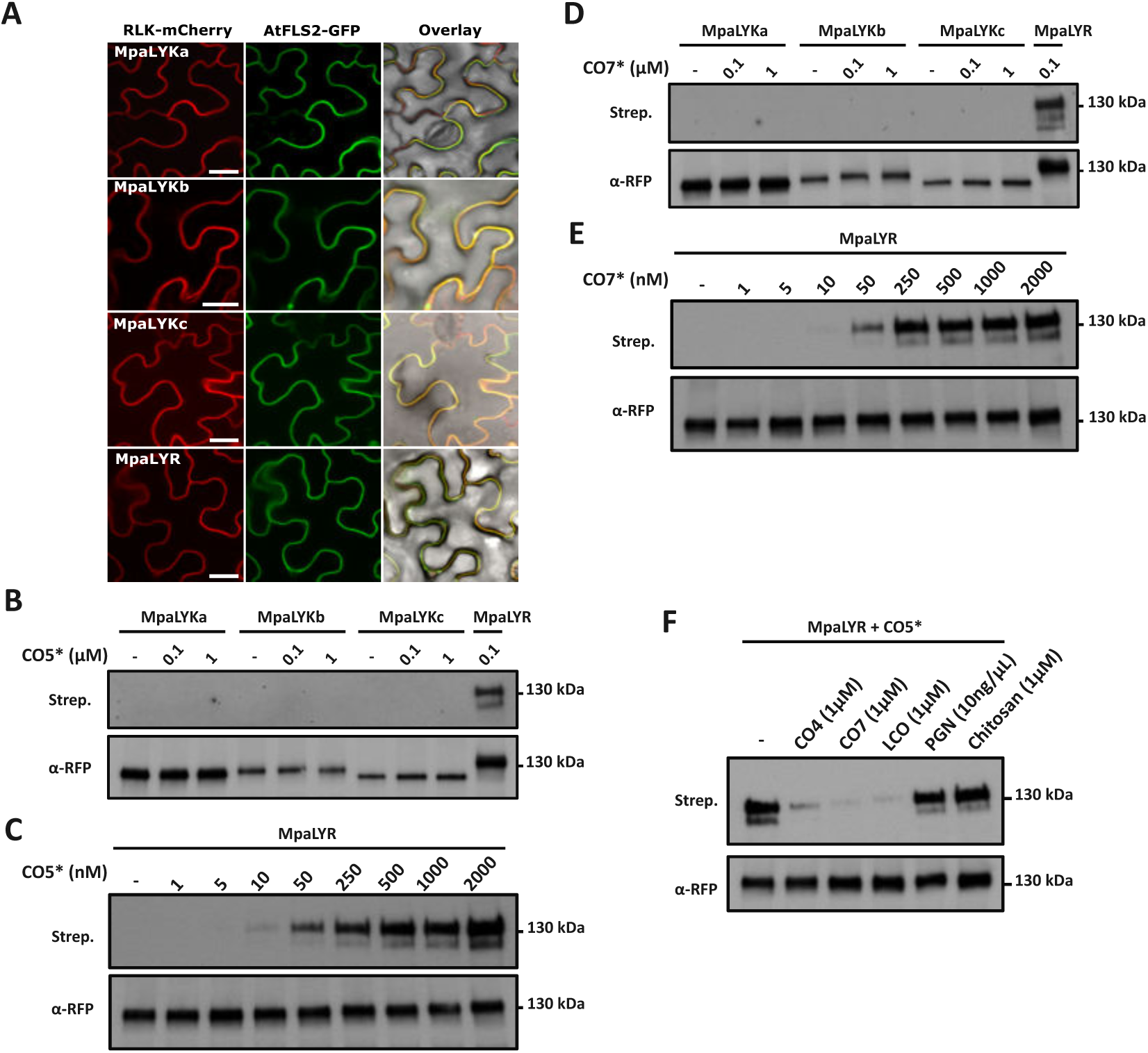
MpaLYR has high affinity for chito- and lipochito-oligosaccharides in *M. paleacea*. **(A)** Confocal micrographs of *Nicotiana benthamiana* epidermal cells transiently co-expressing indicated *M. paleacea* LysM-RLK-mCherry fusions (left panel) and the plasma membrane marker AtFLS2-GFP (middle panel). Overlays including bright field images are shown (left panel). Scale bars = 25 µm. (**B-F**) Microsomal fractions from *N. benthamiana* leaves expressing *M. paleacea* LysM-RLK-mCherry fusions, as indicated, were utilised in binding assays with indicated concentrations of crosslinkable biotinylated chitopentaose (CO5*) or chitoheptaose (CO7*). Ligand bound receptors and total immunopurified receptors were detected using Streptavidin (top panels) or anti-RFP antibodies (bottom panels), respectively. (**F**) CO5*-binding competition assays on MpaLYR using 10-fold higher concentrations of CO4, CO7, Fuc/MeFuc-LCOs, PGN fragments or chitosan fragments relative to CO5*.

### COs and LCOs early signalling is impaired in *lyka* and *lyr* mutants of *Marchantia paleacea*

The function of MpaLYR, but not MpaLYKa, as a receptor for COs and LCOs was in apparent contradiction with the AM phenotypes of the corresponding mutants, with *MpaLYKa* but not *MpaLYR*, being critical for AM establishment. Therefore, we sought to correlate AM establishment and perception of AM fungus-derived signals by determining the signalling abilities of the different *M. palecea* receptor mutants in response to COs and LCOs. Ligand perception events by receptor-like kinases induce within seconds calcium influx into the cytosol in both immune and symbiotic contexts [28]. To use this physiological output, we generated a *M. paleacea* line expressing the cytosolic apoaequorin luminescent marker [29] under the control of the constitutive *MpoEF1ɑ* promoter [30]. Once established and tested for suitable apoaequorin expression, this line served as a homogeneous background to re-create loss-of-function mutants using CRISPR/Cas9, as well as a control line expressing the Cas9 endonuclease alone (AEQ-cas9). Plants were then treated with CO7, CO4 or different LCOs, including Fuc/MeFuc-LCOs, LCO-IV(C18:1) (hereafter NS-LCOs) and LCO-IV(C18:1, S) (hereafter S-LCOs) [27,31].

To determine an effective working concentration for all molecules, we tested a range of concentrations starting at 10^-6^ M (1µM) followed by 10-fold series dilutions (**Fig 5A**). At the concentration of 1 µM, CO7, CO4 and Fuc/MeFuc-LCOs induced an increase in cytosolic calcium concentration in the *M. paleacea* control line that peaked at 5 minutes after application, before reaching back to the baseline at 15 minutes (**Fig 5A**). In contrast, NS-LCOs and S-LCOs failed to elicit a calcium response at this concentration (**Fig 5B**), suggesting variability in the potency of different LCO species. Importantly, water and dimethyl sulfoxide (DMSO) used to prepare stock solutions for CO4 and CO7 or LCOs, respectively, did not trigger any calcium variation in *M. paleacea* (**Fig S5**). Of the three active molecules, CO7 elicited the most pronounced response that gradually decreased in intensity and showed a slight delay when using diluted solutions. A clear response was still visible at 1 nM of CO7, although not statistically different from the lowest tested and poorly active concentration of 0.1 nM (**Fig 5A - insets**). In contrast, CO4 and Fuc/MeFuc-LCOs already failed to elicit a clear response at 0.1 µM, as evidenced by the shapes of the corresponding response curves (**Fig 5A**). In the following experiments, we have therefore chosen the highest concentration of 1 µM for all molecules. To verify whether cytosolic calcium variations occur at later time points, plant responses to CO7, CO4 and Fuc/MeFuc-LCOs were monitored over a one-hour time course. No additional calcium variation was apparent at later time points, suggesting that a 15 minutes time frame was sufficient to capture the entire cytosolic calcium response of *M. paleacea* to these molecules (**Fig 5C**). A five minutes pre-treatment with 1.5 mM lanthanum chloride (La^3+^), a non-specific calcium channel blocker [32], completely abolished the cytosolic calcium variations triggered by CO7, CO4 or Fuc/MeFuc-LCOs (**Fig 5D**). This supports the view that these cytosolic calcium variations are *bona fide* influxes of calcium originating from the apoplast.

**Fig 5.**
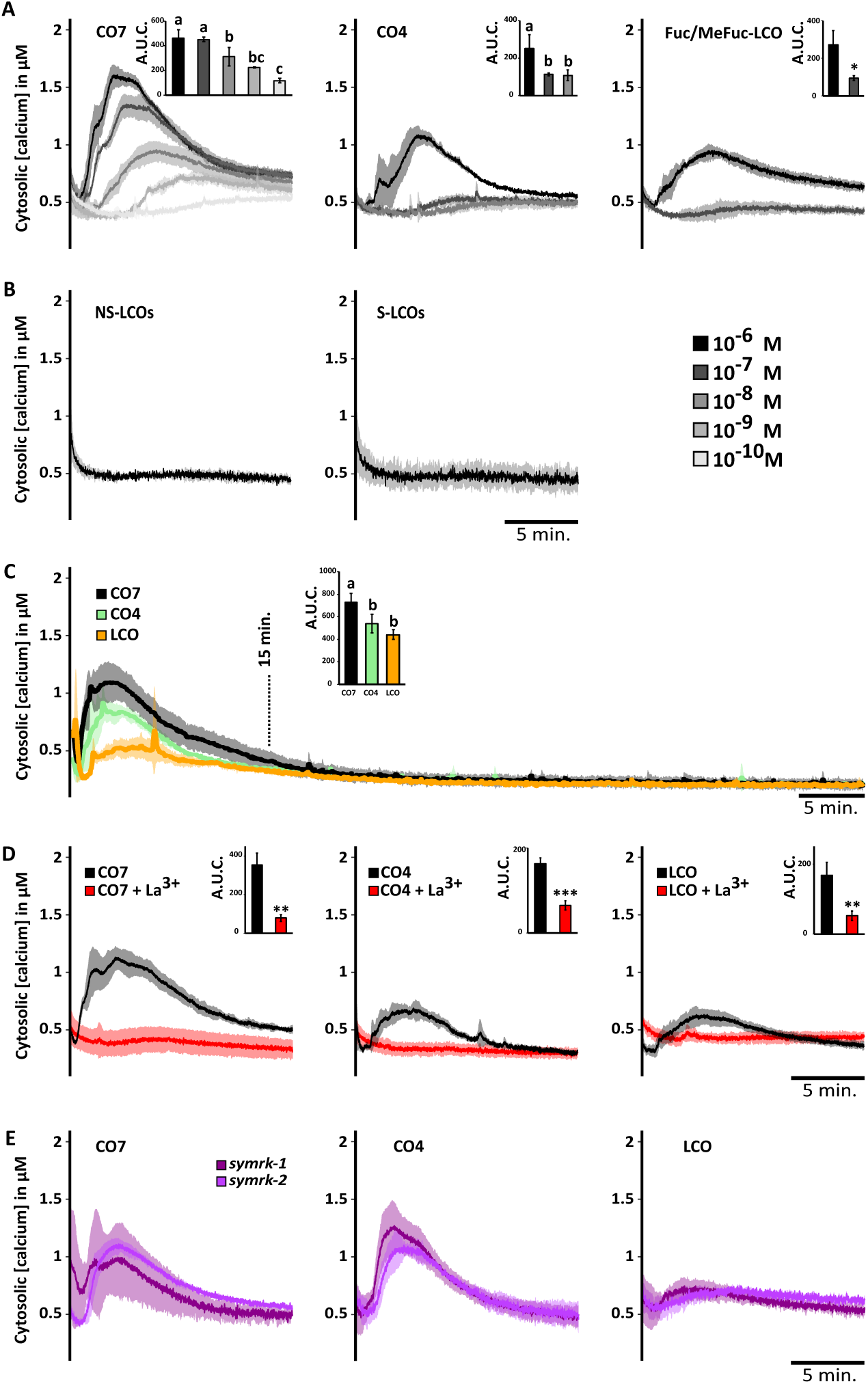
Chito- and lipochito-oligosaccharides induce cytosolic calcium influx in *M. paleacea* independently of the common symbiosis pathway. Cytosolic calcium concentration variations in *M. paleacea* control plants (AEQ-cas9) in response to decreasing concentrations of CO7, CO4 and Fuc/MeFuc-LCOs, as indicated, over a 15 minutes time course. (**B**) Cytosolic calcium concentration variations in *M. paleacea* control plants (AEQ-cas9) in response to 1 µM non-sulphated LCO (NS-LCO) or sulphated LCO (S-LCO), as indicated. (**C**) Cytosolic calcium concentration variations in *M. paleacea* control plants (AEQ-cas9) in response to 1 µM CO7, CO4 or Fuc/MeFuc-LCOs over a one-hour time course. (**D**) Effect of a 5 minutes pre-treatment with 1.5 mM lanthanum chloride on CO7-, CO4- or Fuc/MeFuc-LCO-induced calcium concentration variations. (**E**) Cytosolic calcium concentration variations in two independent *M. paleacea symrk* mutant lines in response to 1 µM CO7, CO4 or Fuc/MeFuc-LCOs. (**A-E**) Each trace represents the mean (line) ± standard deviation (shading) from at least three replicates. (**A-D**) Bar graph insets represent the area under the curves (A.U.C.) corrected for the baseline and different letters indicate significant differences to control conditions inferred by ANOVA followed by Tukey’s HSD post-hoc test (p-value < 0.05) for (A, C) or Student t-test (p-value <0.01**; <0.001***) for (D).

To determine whether this calcium influx triggered by CO7, CO4 and Fuc/MeFuc-LCOs seat upstream or downstream of the CSSP, we analysed the response of two independent *symrk* mutants generated in the aequorin-expressing *M. paleacea* background (**Fig S1 and Table S1**). The duration and amplitude of the calcium influxes measured in AEQ-*symrk-1* and AEQ-*symrk-2* after 1 µM CO7, CO4 or Fuc/MeFuc-LCOs treatments were similar to the control (**Fig 5E**). This result demonstrate that this response is upstream SYMRK and distinct from calcium variations known as calcium spiking, which are strictly dependent on CSSP activation. Taken together, our results show that CO7, CO4 and Fuc/MeFuc-LCOs induce an early calcium influx in *M. paleacea* and that this physiological response depends on a perception module that is independent of SYMRK.

Having established some characteristics of this CO7-, CO4- and Fuc/MeFuc-LCOs-induced calcium influx in *M. paleacea*, we further investigated the function of each LysM-RLK in generating this response. For that, we recovered two independent CRISPR/cas9-edited *M. paleacea* lines for each receptor encoding gene in the aequorin-expressing background (**see. Table S1**). Next, we treated these plants with 1 µM CO7, CO4 or Fuc/MeFuc-LCOs and monitored their calcium influx response in comparison to the AEQ-cas9 control line (**Fig 6**). The duration and amplitude of the calcium influxes obtained in AEQ-*lykb* and AEQ-*lykc* lines were comparable to the control for all three elicitors, whereas AEQ-*lyka* and AEQ-*lyr* lines failed to respond to all treatments (**Fig 6**). Quantitative analysis of the area under the curves and analysis of variance confirmed the similarity between control, AEQ-*lykb* and AEQ-*lykc* and the differences between control and AEQ-*lyka* or AEQ-*lyr* in response to all elicitors (**Fig 6 – bar graph insets**). To rule out the possibility that *lyka* and *lyr* mutations have a pleotropic effect on the plants ability to mount a calcium response, we treated all genotypes with hydrogen peroxide (H_2_O_2_) at a final concentration of 1 mM (**Fig S6**). All aequorin-expressing LysM-RLK mutants responded with an immediate and robust calcium influx and quantitative analysis of the area under the curves and analysis of variance confirmed that there was no difference between control, AEQ-*lyka* and AEQ-*lyr* mutants in response to H_2_O_2_. In addition, the AM fungi colonisation status of AEQ-*lyka* and AEQ-*lyr*, the two non-responsive mutants to COs and Fuc/MeFuc-LCOs, was evaluated after *R. irregularis* inoculation and histological observations confirmed that AEQ-*lyka* was AM-defective (**Fig S7**). Conversely, as previously described (**Figs 2 and 3, S2 and S3**), arbuscules were clearly visible in AEQ-cas9 and AEQ-*lyr* plants with no differences in the size of the exclusion zone (**Fig S7**).

**Fig 6.**
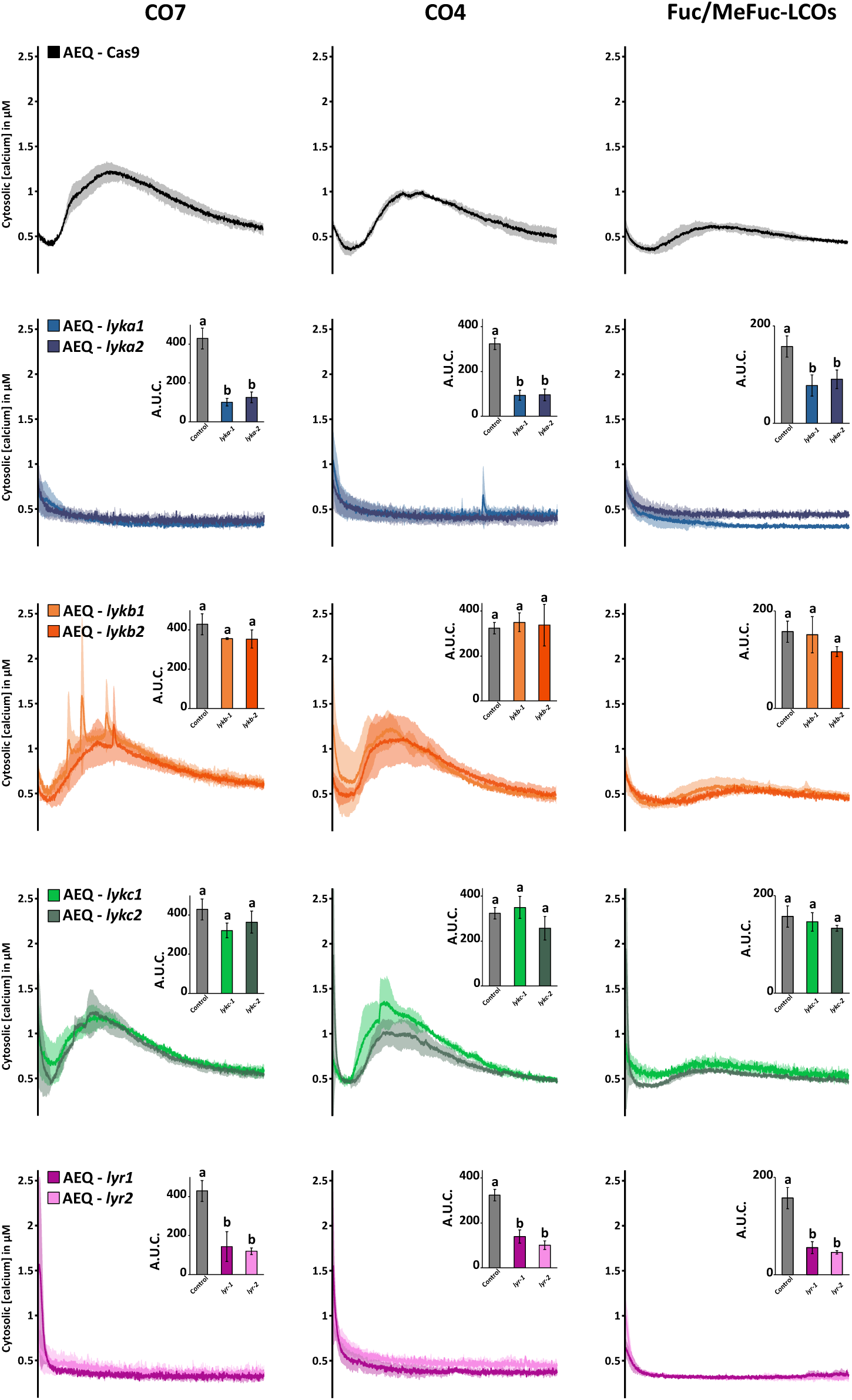
Chito- and lipochito-oligosaccharides-induced calcium influx response in *M. paleacea* depend on *MpaLYKa* and *MpaLYR*. Cytosolic calcium concentration variations in *M. paleacea* control (AEQ-cas9) or two independent mutant lines for each *LysM-RLK* of *M. paleacea*, as indicated. Plants were treated with 1 µM CO7 (left), 1 µM CO4 (middle) or 1 µM Fuc/MeFuc-LCOs (right). Each trace represents the mean (line) ± standard deviation (shading) from at least three replicates over a time course of 15 min. Bar graph insets represent the area under the curves (A.U.C.) corrected for the baseline and different letters indicate significant differences versus control inferred by ANOVA and Tukey’s HSD post-hoc test (p-value < 0.05).

Collectively, these results demonstrate that MpaLYKa and MpaLYR are both required for the perception of chitin-based molecules and their early signalling in *M. paleacea*.

### CO7-induced transcriptomic changes are similarly affected in *lyka* and *lyr* mutants

The perception of non-self and the activation of corresponding signalling pathways in plants often culminate in adaptive transcriptomics reprogramming. Having established that early signalling in response to fungal molecules was blocked in both the *lyka* and *lyr* mutants, we then tested whether a later response such as gene induction was also affected in these mutants. First, we performed genome-wide transcriptomics of *M. paleacea* empty vector control plants after a one-hour treatment with 1 µM of CO7 or CO4. When compared to treatment with water only, CO7 and CO4 induced a statistically significant differential expression for 444 and 108 genes (p-adjusted value ≤ 0.05), respectively (**Table S2**). Of the differentially expressed genes (DEGs) identified in response to CO7, 195 were up-regulated with a log2 fold induction (log2FC) greater than 1, while 16 were down-regulated with a log2FC less than −1 (**Fig 7A**). In response to CO4, the application of identical log2FC cut-off values identified 45 genes that were up-regulated and two genes that were down-regulated (**Table S2**). Interestingly, 87% (94/108) of DEGs identified in response to CO4 are shared with CO7 (**Fig S8A**) and a direct comparison of the log2FC values in response to both treatments for the 108 CO4-responsive genes resulted in a Pearson’s correlation coefficient of 0.91 (**Fig S8B**). The overlap and strong correlation between DEGs groups suggest that the transcriptomic response to CO4 represents a subset of the response to CO7 in *M. paleacea*. Furthermore, CO7 appears to be a more potent elicitor of gene reprogramming in this species, as evidenced by the fourfold higher number of identified DEGs in response to this molecule.

**Fig 7.**
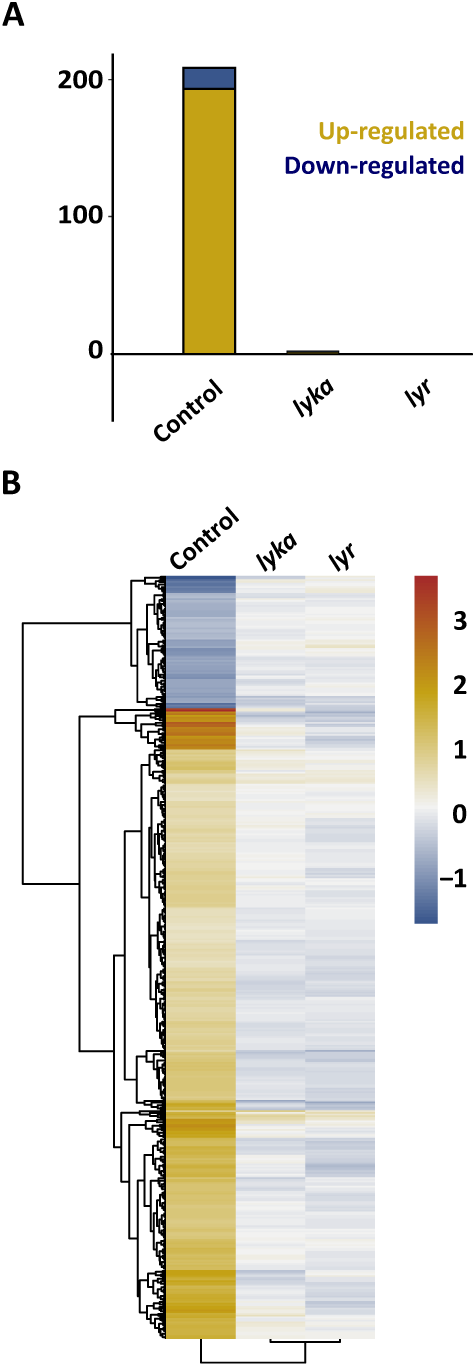
*MpaLYKa* and *MpaLYR* are essential for gene induction in response to CO7. Number of significantly upregulated or downregulated genes (yellow or blue, respectively) in CO7-treated plants for one-hour (control, *lyka* and *lyr*) relative to non-treated plants. DEGs are statistically significant (p-value adjusted < 0.05 and 1log2FC1 ≥ 1). (**B**) Heatmap of the expression pattern in *lyka* and *lyr* mutants compared with the control in plants treated with 1µM CO7. Displayed fold changes are statistically significant for the control (p-value adjusted < 0.05).

In order to gain insight into the most represented molecular functions, an InterPro domain enrichment analysis was performed on the up- and down-regulated genes in response to CO7 (**Fig S9 and Table S3**). A total of 18 InterPro terms were found to be significantly enriched among the subset of CO7-up-regulated genes. These included the terms “Leucine-rich repeat domain superfamily (IPR032675),” “Protein kinase domain” (IPR000719) and “WRKY domain superfamily” (IPR036576). The InterPro domain enrichment analysis using the set of 45 up-regulated genes after CO4 treatment yielded similar results, albeit with a lower number of significantly enriched terms (15 versus 18) (**Fig S10 and Table S3**). Taken together, these results suggest that CO perception in *M. paleacea* induces genes involved in cell-surface signalling and transcription. Conversely, the set of 16 down-regulated genes following CO7 treatment was significantly enriched in genes with functions related to photosynthesis and carbon fixation (**Fig S9B**), suggesting a potential growth inhibition trade-off after CO7 perception. The two down-regulated genes after CO4 treatment were not sufficient to perform a similar enrichment analysis.

In view of the overlap between the CO7 and CO4 transcriptomic responses observed in the control plants, we assessed the response of the *lyka* and *lyr* mutants to CO7 only. In the *lyka* mutant, CO7 treatment resulted in a statistically significant differential expression for 47 genes, with only two genes being up-regulated and two genes being down-regulated when the previously described log2FC cut-off values were applied (**Fig 7A**). Moreover, none of these four up- or down-regulated genes were present in the list of DEGs identified in the control plants (**Table S2**), suggesting that they may represent noise rather than a true signal. In the *lyr* mutant, CO7 did not elicit any response in comparison to the water treatment, as none of the two DEGs identified in this mutant showed a log2FC above the cut-off values (**Fig 7A and Table S2**). A comparison across genotypes of the log2FC values from the DEGs identified in the CO7-treated control plants further highlights the lack of response in the *lyka* and *lyr* mutants (**Fig 7B**). Altogether, these results indicate that the *lyka* and *lyr* mutants of *M. paleacea* are equally unable to induce gene expression in response to CO7.

## Discussion

While single mutants in various *LYKs* in angiosperms have been observed to exhibit delayed or reduced colonization rates [10,33–37], none is completely defective for AM. In contrast, this work describes a single LYK mutant that displays an AM phenotype indistinguishable from those observed in mutants of the CSSP in angiosperms [4,5,38–43] or *M. paleacea* [8,9]. The absence of AM in *lyka* is consistent with the AM-defective phenotypes described in the *M. truncatula lyk8/lyk9* [14] and in the *L. japonicus lys6/lys7* double mutants [13]. This result obtained in a bryophyte species, combined with the phenotypes of the double mutants in legumes, corroborate the ancestral role of the LYK-I subclade in AM and substantiate the hypothesis that perception of fungal-derived molecule(s) by LysM-RLK is a prerequisite for AM establishment. Conversely, the loss of function of either *MpaLYKb* or *MpaLYKc* did not affect the establishment of AM. While no symbiotic function has yet been described for members of the LYK-III subclade (*MpaLYKc*), the co-orthologues of *MpaLYKb* (LYK-II subclade) in *L. japonicus*, *LjEPR3* and *LjEPR3a*, have been proposed to play regulatory functions during the accommodation of microbes [44,45]. *LYK-11* genes are also upregulated during AM in a range of plant species, including monocots [46], dicots, and bryophytes (for review, see Buendia et al., 2018). Interestingly, they are conserved in plants that form endosymbioses and are otherwise lost, a phylogenetic pattern that has been observed for genes of the CSSP [23]. These two lines of evidence support the existence of a conserved function for members of the LYK-II subclade in endosymbioses. The persistence of AM formation in the *LYK-11* mutants of *M. paleacea, L. japonicus* and rice suggests however that this function might come in complement to other signalling cues.

In addition to providing evidence supporting the involvement of MpaLYKa in AM, we demonstrate through measurement of cytosolic calcium influx that MpaLYKa is essential for both COs and LCOs perception. This calcium influx following CO and LCO perception in *M. paleacea* is independent of *SYMRK*, suggesting that it is distinct from the calcium spiking response observed in and around the nucleus, which strictly requires activation of the CSSP [47,48]. This observation is consistent with research in legumes, demonstrating the existence of an early calcium influx independent of the CSSP in response to Nod-factors [49] or CO8 and CO4 [50].

Perception of COs/LCOs in plants often involves heterocomplexes between LysM-RLKs of the LYK-I and LYR subclades [10], raising the possibility of an interaction between of MpaLYKa and MpaLYR for CO/LCO signalling in *M. paleacea*. In angiosperms, the LYR subgroup has expanded, giving rise, for example, to four and eleven paralogs in *Brachypodium distachyon* and *M. truncatula*, respectively [10]. The potential function of LYRs during AM has been extensively studied and single mutants have been tested for their symbiotic abilities in *M. truncatula* (*nfp* and *lyr4*) [51,52], *L. japonicus* (*Ljlys11*) [53], *Parasponia andersonii* (*Pannfp1*, *Pannfp2*) [37], tomato (*sllyk10*) [36], Petunia (*Phlyk10*) [36], barley (*rlk2* and *rlk10*) [39], *B. distachyon* (*bdlyr1* and *bdlyr2*) [54] and rice (*Osnfr5*) [55]. In addition, the double mutants *M. truncatula nfp/lyr4* [51] and *nfp/lyr1*, *L. japonicus lys11/nfr5* [53], *B. distachyon bdlyr1/bdlyr2* [54] and barley *rlk2/rlk10* [39] were tested. Most of the single mutants displayed wild-type level of colonisation by AM fungi, with the exception of tomato and petunia single mutants, and even double mutants were either not or only partially affected in their AM fungi colonisation levels. One hypothesis to explain these mixed results is the occurrence of significant variation in the number of LysM-RLKs between these species and possible genetic redundancy or variable contributions from the different paralogs in a species-specific manner. In *M. paleacea*, only one member of the *LYR* subgroup is found and AM-phenotyping of the *lyr* loss-of-function mutants demonstrates the dispensable role of this receptor for AM establishment in this species.

Interestingly, MpaLYR is able to bind short- and long-chain COs with high affinity. The ability of MpaLYR to bind LCO molecules is probably due to the presence of the CO moiety of the LCO rather than to a specific binding of the lipid moiety. The biochemical properties of MpaLYR are thus reminiscent of those described for angiosperm LYR-IB receptors [10]. This supports an evolutionary scenario in which receptors with specificity for LCO, such as members of the LYR-IA [12,36,56] and LYR-IIIA [57,58] subclades, and receptors with specificity for long-chain versus short-chain COs, such as members of the LYR-IIIC subclade [14,59], have emerged after the LYR expansion in angiosperms. This could have led to a fine-tuning of the perception of symbionts and pathogens.

In contrast to MpaLYR, no binding activity towards chitin-based molecules could be detected in any of the MpaLYK receptors. A similar observation has been reported in *M. truncatula* concerning the ligand-binding abilities of LYR and LYK proteins [14]. However, both MpaLYR and MpaLYKa are essential for the perception of chitinaceous compounds in *M. paleacea*. This suggests the existence of MpaLYR/MpaLYK heterocomplexes for their perception, with a ligand-binding property conferred by MpaLYR. Indeed, we have demonstrated that *lyr* and *lyka* mutants were incapable of producing a calcium influx in response to CO7, CO4 or LCOs. Furthermore, CO7-dependent gene reprogramming was entirely abolished in *lyr*, as it was in *lyka*. These results mirror the critical role of both MpoLYK1 and MpoLYR in CO7-triggered ROS production in the non-AM-host species *Marchantia polymorpha* [20].

The identification of bacterial lipo-chitooligosaccharides (Nod-LCOs) that are essential for the nitrogen-fixing symbiosis in legumes has led to the long-held hypothesis that functional analogues are produced by AM fungi. The discoveries of AM fungi-produced LCOs and short-chain COs with symbiotic activity in plants [52,60] provided compelling evidence to support this hypothesis. It has been proposed that LCOs and short-chain COs represent a blend of symbiotic signals produced by AM fungi to initiate symbiotic responses [52,60,61]. However, it remains unclear whether the perception of these molecules by the host is a prerequisite for AM establishment. Interestingly, other compounds, such as CO8, can induce similar molecular and cellular responses [14]. Furthermore, large-scale analyses of exudates from sixty fungal species revealed that neither short-chain COs nor LCOs are specific to AM fungi [27,62], raising questions about how plants can discriminate between beneficial and detrimental organisms based on this class of compounds alone. Our work in *M. paleacea* suggests that AM establishment can be uncoupled from COs and LCOs perception events and that MpaLYKa-dependent perception of other type of symbiont-derived signal(s) occurs. The suite of *M. paleacea* mutants developed here offer the opportunity to identify these elusive signal(s) in the future.

The study of mutants in bryophyte species which have lost the ability to form symbiosis allowed discovering the ancestral function of LysM-RLK in plant immunity over the last decade [20,63]. Our work extends this knowledge and proposes that LysM-RLK have been maintained in land plant genomes for almost half a billion years not only to perceive pathogens but as readers of the biotic environment, both for immunity and symbiosis.

## Materials and methods

### Phylogeny

To reconstruct the phylogeny of LysM-RLKs, a set of 102 protein sequences composed of 8 Marchantia genes and the set described by Buendia et al. (2018) supplemented with the *Lotus japonicus* EPR3 and EPR3a sequences [45] was aligned using MUSCLE (v3.8). The phylogeny was reconstructed using IQ-TREE v1.6.12 (http://iqtree.cibiv.univie.ac.at/) with the VT+F+I+G4 model and support was provided with 1,000 ultra-fast bootstrap replicates [64–66]. The unrooted tree visualisation was generated using iTOL [67] and adapted for presentation using InkScape.

### Plant material, growth conditions and transformation

*Marchantia paleacea ssp. paleacea* wild-type plants were grown axenically *in vitro* from sterile gemmae on ½ strength Gamborg B5 (G5768, Sigma) medium supplemented with 1.4 % agar (1330, Euromedex) for 3 to 4 weeks in 16/8h photoperiod at 22°C/20°C. For transformation, ∼20 plantlets were blended 15 seconds in a sterile 250mL stainless steel bowl (Waring, USA) in 10mL of OM51C medium [KNO_3_ (2 g.L^-1^), NH_4_NO_3_ (0.4 g.L^-1^), MgSO_4_ 7H_2_O (0.37 g.L^-1^), CaCl_2_ 2H_2_O (0.3 g.L^-1^), KH_2_PO_4_ (0.275 g.L^-1^), L-glutamine (0.3 g.L^-1^), casamino-acids (1 g.L^-1^), Na_2_MoO_4_ 2H2O (0.25 mg.L^-1^), CuSO_4_ 5H_2_O (0.025 mg.L^-1^), CoCl_2_ 6H_2_O (0.025 mg.L^-1^), ZnSO_4_ 7H2O (2 mg.L^-1^), MnSO_4_ H2O (10 mg.L^-1^), H_3_BO_3_ (3 mg.L^-1^), KI (0.75mg.L^-1^), EDTA ferric sodium (36.7 mg.L^-1^), myo-inositol (100 mg.L^-1^), nicotinic acid (1 mg.L^-1^), pyridoxine HCL (1 mg.L^-1^), thiamine HCL (10mg.L^-1^)]. The blended plant tissues were transferred to 250mL erlenmeyers containing 50 mL of OM51C and kept at 22°C/20°C, 16/8h photoperiod, 200 RPM for 3 days. Co-cultures with *A. tumefaciens* (GV3101) containing constructs were initiated by adding 1mL of saturated bacterial liquid culture (DO = 0,5, 50 mL final), acetosyringone (100 µM final) and glutamine (0,5 g.L^-1^ final). After 3 days, plant fragments were washed by decantation 3 times with water and plated on ½ Gamborg containing 200 mg.L^-1^ amoxicillin (Levmentin, Laboratoires Delbert, Fr.) and 10 mg.L^-1^ Hygromycin B (3250, Euromedex) or 50 nM Chlorsulfuron (34322-100MG Sigma). Plant material is available in **Table S5**.

For AM experiments, *Marchantia paleacea ssp. paleacea* control and mutants were grown on a zeolite substrate (50% fraction 1.0-2,5mm, 50% fraction 0,5-1.0-mm, Symbiom) in 7×7×8 cm pots with a density of five thalli per pot. Pots were watered once a week with Long Ashton medium containing 7.5 μM of phosphate [68] and grown with a 16/8h photoperiod at 22°C/20°C for at least 2 weeks prior inoculation with AM fungi. For calcium measurement, similar conditions were applied with the exception of a higher plant density per pot and a growth duration of 4 weeks or plants were grown axenically *in vitro* on 0,4% phytagel diluted in Long Ashton medium containing 7.5 μM of phosphate. For RNAseq experiments, plants were grown axenically *in vitro* on 0,4% phytagel diluted in Long Ashton medium containing 7.5 μM of phosphate for a duration of 4 weeks.

### Cloning

The Golden Gate modular cloning system was used to prepare the plasmids as described in *Schiessl et al* [69]. Level 1 and Level 2 empty vectors used in this study are listed in the **Table S5** and held for distribution in the ENSA project core collection (https://www.ensa.ac.uk/). Other Level 0, Level 1 and Level 2 vectors containing constructs are listed in the **Table S5**. Coding sequences were either domesticated in-house by PCR in the pUPD2 vector or synthesised and cloned into pMS (GeneArt, Thermo Fisher Scientific, Waltham, USA).

### Transient expression assays in *Nicotiana benthamiana*

*Agrobacterium tumefaciens* (GV3101) cultures carrying vectors of interest were grown under agitation overnight at 28°C in liquid LB medium supplemented with adequate antibiotics. Cultures were washed twice by centrifugation at 8000g and re-suspended in water supplemented with 50µM acetosyringone. Bacteria concentrations were adjusted to OD_600_=0.2 per construct prior infiltration of the abaxial side of two months old *N. benthamiana* leaves using a 1 mL needle-less syringe.

### Microscopy

For microscopy analyses, mCherry and GFP signals were sequentially acquired 48 hours after *N. benthamiana* transfection using a Leica SP8 confocal microscope mounted with a FLUOTAR VISIR 25x/0.95NA water immersion objective (Leica). GFP was excited at 488 nm and detected from 500 to 535 nm; monomeric Cherry was excited at 552 nm and detected from 560 to 610 nm. Images were processed using LasX (Leica).

### Binding assays

Preparation of microsomal fraction was performed as previously described by Girardin et al., 2019 [36]. CO5-biotin (CO5*) and CO7-biotin (CO7*) were synthesised as detailed by Cullimore et al., 2023 [56] and Ding et al., 2024 [54] respectively. Binding assays were conducted using CO5* or CO7* and competitors, mixed with the appropriate microsomal fraction in binding buffer (25 mM NaCacodylate pH 6, 1 mM MgCl2, 1 mM CaCl2, 250 mM sucrose and protease inhibitors). Samples were incubated for 30 min on ice and subsequently centrifuged for 30min at 31 000g and 4°C. Pellets were resuspended in IP buffer (25 mM Tris-HCl pH7.5, 150 mM NaCl, 10% glycerol, supplemented with protease inhibitors). Proteins were solubilized in IP buffer supplemented with 0.2% DDM for 1h at 4°C and centrifuged during 30 min at 31 000g and 4°C. The supernatant was retrieved, and mCherry-tagged receptors were immunoprecipitated using RFP-trap magnetic agarose beads (ChromoTek). Beads were washed twice with IP buffer, and proteins were extracted in Laemmli buffer. Proteins were separated by SDS-PAGE and transferred onto a nitrocellulose membrane using the Trans-Blot system (Bio-rad) according to the manufacter’s instructions. Membranes were blocked for 1h with 5% fat-free milk in Tris-buffer Saline (TBS) supplemented with 0.1% Tween-20 (TBS-T). Membranes were incubated for 1h with the appropriate antibodies: rabbit αRFP [70] (1:5000), αRabbit-Hrp (Millipore, 1:20000), and streptavidin-Hrp (Invitrogen, 1:3000). Hrp activity was detected using Clarity Western ECL substrate (Bio-Rad) and observed with a ChemiDoc imager (Bio-Rad).

### Generation of CRI5PR mutants in *M. paleacea ssp. paleacea*

Constructs containing the *A. thaliana* codon-optimised SpCas9 [71] under the control of *MpoEF1ɑ* promoter and single guide RNAs (**Table S1**) under the *M. polymorpha* or *M. paleacea* U6 promoters were introduced in wild-type *M. paleacea.* Independent transformants for *lyka, lykb, lykc,* and *lyr* were selected for AM phenotyping. A *M. paleacea* line expressing the calcium reporter aequorin (AEQ) under the control of *MpoEF1ɑ* promoter was re-transformed with the CRISPR/Cas9 vectors, modified to contain a secondary selection marker (ChlorsulfuronR). Two independent mutant lines per receptor were used for calcium influx assays.

### Mycorrhization tests

Each pot containing 5 plants was inoculated with ∼1,000 sterile spores of *Rhizophagus irregularis* DAOM 197198 (Agronutrition). For the nursing experiment, thalli of empty-vector control and *lyka* were transplanted in proximity to wild-type plants inoculated with *R. irregularis* spores more than 10 weeks before. Six- or ten-weeks post-inoculation (for nursing), thalli were cleared of chlorophyll using ethanol 100% for 1 day before storage in an EDTA (0.5 mM). Cleared thalli were scanned with an EPSON 11000XL and the distance between apical notches and the colonisation zone (naturally coloured with a purple/brown pigment) were scored on using ImageJ. Large-scale mycorrhiza assays were run independently twice.

For imaging, cleared thalli of control and mutants were embedded in 6% agarose and 100 µm thick transversal and longitudinal sections were prepared using a Leica VT1000s vibratome. Sections were incubated overnight in 10% K-OH (w/v) prior to three washes in PBS. Fungal membranes were stained overnight using 1 µg.mL^-1^ WGA-Alexa Fluor 488 (Sigma) diluted in PBS. Pictures were taken with a Zeiss AxioZoom V16 binocular using a PlanApo Z 0.5X. objective and the ZEN software suite with similar settings. Alexa Fluor was excited between 450 and 490 nm and emission collected between 500 and 550 nm.

### Calcium influx measurements

Three to four weeks old plants were submerged in 2.5 μM coelenterazine-h (Interchim) diluted in water for a minimum of 16 hours at room temperature and in the dark. Samples were then transferred in a Berthold Sirius luminometer before treatments with a 250 µL aqueous solution of 1 µM CO7 dilutions, 1 µM CO4 (Elicityl, Crolles, France) or a 1 µM mixture of fucosyl/methylfucosyl LCO-V (C18:1, Fuc/MeFuc), 1 µM non-sulphated LCOs and 1 µM sulphated LCOs. Luminescence was continuously recorded for 15 min using a 1 sec interval. An equal volume of 2X lysis buffer [20% ETOH, 10mM CaCl2, 2% NP-40] was injected to discharge the total luminescence left and light was collected for an additional 5 min. Calcium concentrations were calibrated as previously done [72]. Graphs were prepared using RStudio and the ggplot2 package by plotting signals from 10 to 900 seconds after treatment, prior to assembly using the InkScape software. Areas under the curve from 10 to 900 seconds were calculated minus the baseline value, defined as the lowest value present in each trace. A minimum of three traces for each experiment were recorded.

### RNA extraction & RNA sequencing

Gemmae from control Cas9, *lyka-1* and *lyr-1* were surface sterilized with a nylon cell strainer 40µm (Falcon) in a bleach solution (∼0.27%) for 1 minute and washed in water twice before plating on ½ strength Gamborg B5 media and grown with a 16/8h photoperiod at 22°C/20°C. Plants were treated or not with 1 µM elicitors for 1 hour before being frozen in liquid nitrogen. Tissues were ground with mortar and pestle using liquid nitrogen prior total RNAs extraction using the RNA Plant and Fungi NucleoSpin™ (Macherey-Nagel™). DNA was eliminated using the Macherey-Nagel™ Kit rDNase. RNAs quality and quantity were determined by Nanodrop. Total RNAs were sent for sequencing to Genewiz/Azenta (Leipzig, Germany). Illumina libraries were prepared with the NEBnext ultra II RNA directional kit and sequenced on a NovaSeq platform. The samples were sequenced using a 2×150 Pair-End reads configuration.

### RNAseq analysis

RNASeq raw reads from all described conditions were mapped against the genome of *M. paleacea* [17] and counted using the Nextflow v23.10.0 [73] pipeline NF-CORE/RNASeq v3.14 [74] with the options star_salmon to align and quantify reads, as well as ‘-nextseq 30 -length 50’ as extra parameters of TrimGalore v0.6.7 [75] to remove reads with quality lower than 30 or a length lower than 50bp. Ribosomal RNA were also removed through the option “-remove_ribo” using SortMeRNA v4.3.4 [76]. The pipeline was run under the GenoToul configuration (available here: https://github.com/nfcore/configs/blob/master/docs/genotoul.md) and used the following softwares and languages: bedtools v2.30.0 [77], R v4.0.3, v4.1.1 and v4.2.1 [78], DESEQ2 v1.28.0 [79]; dupradar v1.28.0 [80], fastqc v 0.12.1 [81], fq v0.9.1 (https://github.com/stjude-rust-labs/fq), gffread v0.12.1 [82], perl v5.26.2 [83], python v3.9.5 [84], rsem v1.3.1 [85], STAR v2.7.10a [86], picard v3.0.0 [87], qualimap v2.3 [88], rseqc v5.02 [89], salmon v1.10.1 [90], summarizedExperiment v1.24.0 [91], samtools v1.16.1 [92], stringtie v2.2.1 [93], tximeta v1.12.0 [94], UCSC v377 and v445 (https://github.com/ucscGenomeBrowser/kent). The R package DESeq2 v1.42.1 was used with default parameters for the differential gene expression analysis. Venn diagram was constructed using the https://jvenn.toulouse.inrae.fr/ webserver with size modification in Inkscape. Enriched IPR terms were determined using the R package clusterProfiler v4.12.3 [95], the DEGs with a log2FC threshold of 111 as foreground and genes subjected DEG analysis (i.e. genes that were retained after removing the low counts) as background. A template script is available at https://gist.github.com/LRSV-MMbengue/beb55a608d6053c9aaa9257993c6bf6b

### Statistical analyses

To evaluate differences versus control, student t-test or ANOVA followed by Tukey’s HSD as post-hoc test was performed using R.

## Supporting information

**Fig S1.**
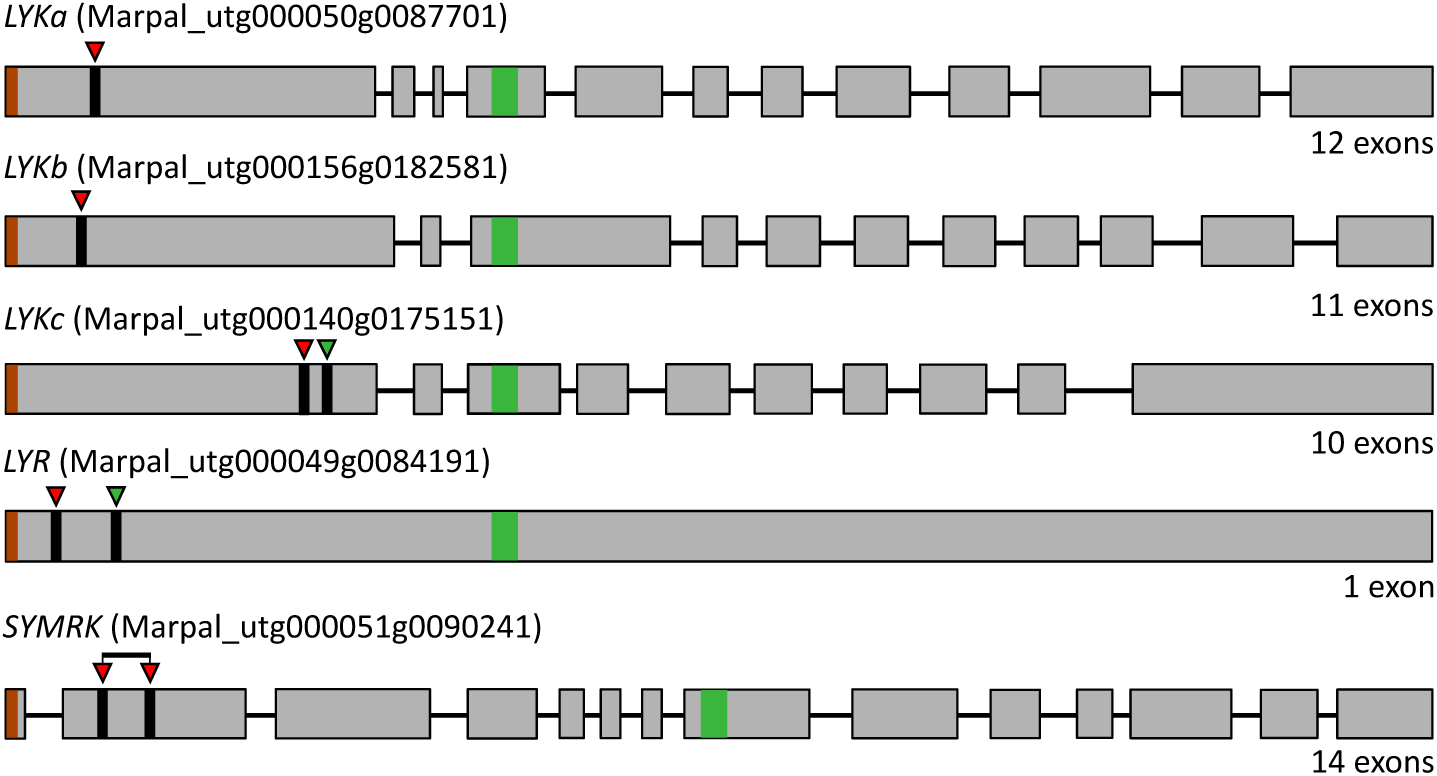
Protospacers positions for generating loss-of-function RLK mutants in *M. paleacea*. Gene structure representation of the four *M. paleacea LysM-RLKs* and *SYMRK:* exons (grey boxes), introns (black line), signal peptides and transmembrane domain coding sequences are highlighted in brown and green, respectively. Arrowheads point at protospacers positions used for generating CRISPR/Cas9 loss-of-function mutants. For *SYMRK*, a polycistronic RNA construct was designed to generate two guide RNAs.

**Fig S2.**
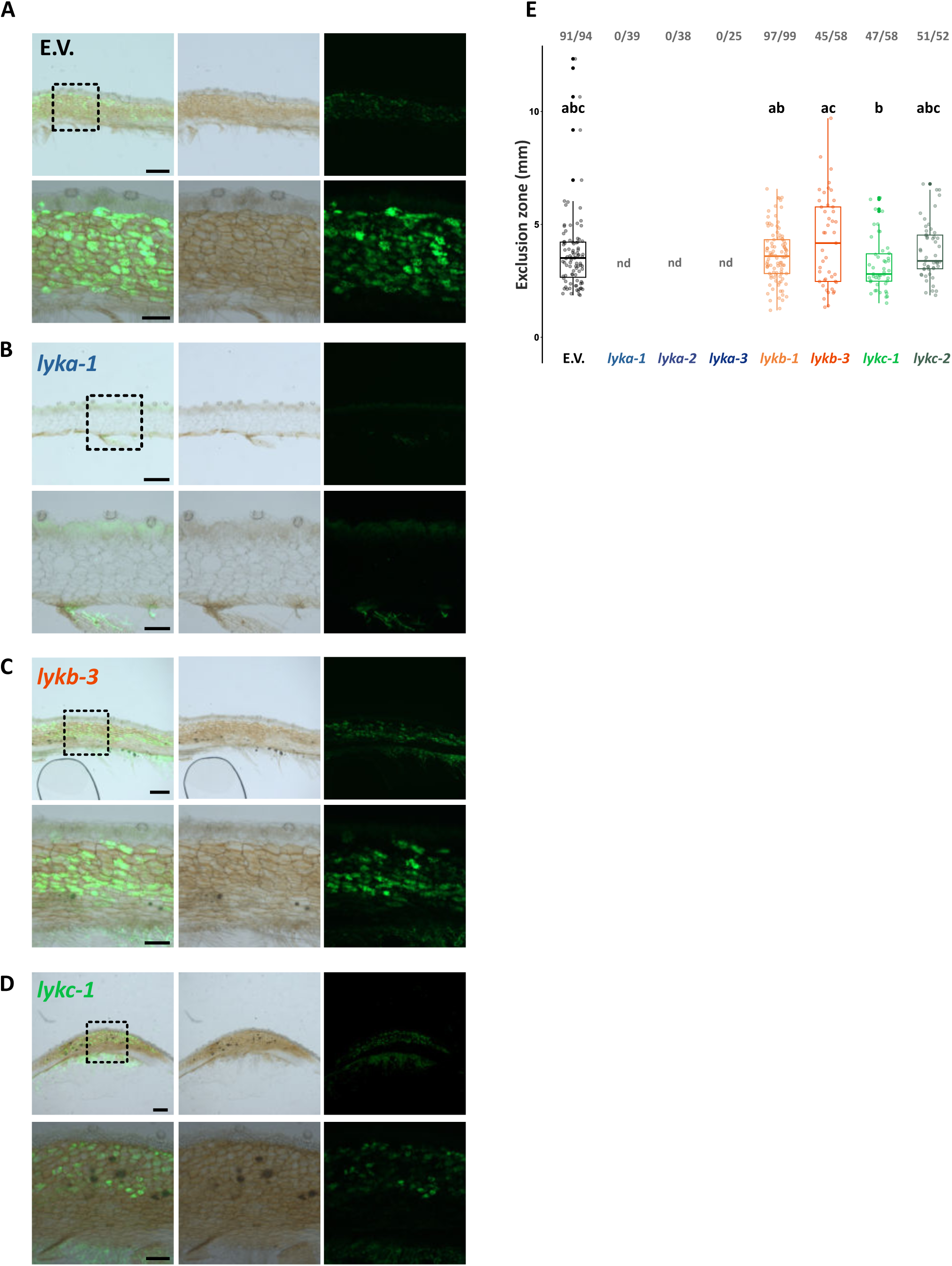
Longitudinal sections of *M. paleacea lyk* mutants and exclusion zone measurements. **(A)** Representative images of longitudinal sections of *M. paleacea* control (E.V.) or *lyk* mutants, as indicated, six weeks after inoculation with *R. irregularis*. Left panels are overlays of bright field (middle panels) and fluorescent (right panels) images. Wheat germ agglutinin (WGA) coupled to Alexa Fluor 488 was used to detect fungal structures. Dashed line delimited insets are enlarged underneath the original images. Scale bars are 500 µm and 200 µm for insets. (**B**) Quantitative analysis of the exclusion zone length on mycorrhizal plants for control (E.V.) and loss-of-function *lyk* mutants. For each genotype, fractions in bold grey represent mycorrhizal thalli over total thalli assessed. Different letters indicate differences to control inferred by ANOVA followed by Tukey’s HSD post-hoc test (p-value < 0.05). Control (E.V.) values are shared with **Fig S3** and statistical analysis was performed on the whole dataset. “nd” stands for not determined.

**Fig S3.**
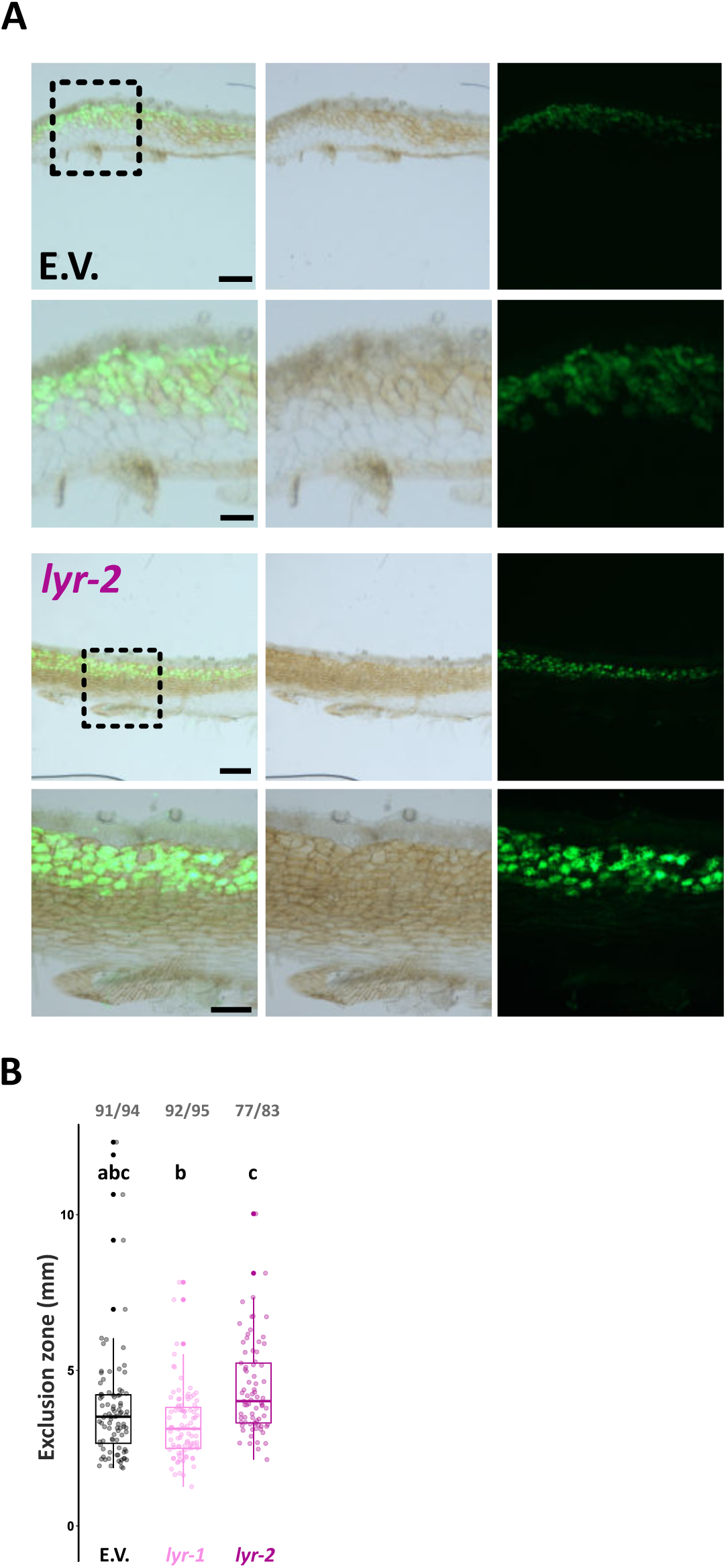
Longitudinal sections of *M. paleacea lyr* mutant and exclusions zones measurements. **(A)** Representative images of longitudinal sections of *M. paleacea* control (E.V.) or the *lyr-2* mutants six weeks after inoculation with *R. irregularis*. Left panels are overlays of bright field (middle panels) and fluorescent (right panels) images. Wheat germ agglutinin (WGA) coupled to Alexa Fluor 488 was used to detect fungal structures. Dashed line delimited insets are enlarged underneath the original images. Scale bars are 500 µm and 200 µm for insets. (**B**) Quantitative analysis of the exclusion zone length on mycorrhizal plants for control (E.V.) and two independent *lyr* mutants. For each genotype, fractions in bold grey represent mycorrhizal thalli over total thalli assessed. Different letters indicate differences to control inferred by ANOVA followed by Tukey’s HSD post-hoc test (p-value < 0.05). Control (E.V.) values are shared with **Fig S2** and statistical analysis was performed on the whole dataset.

**Fig S4.**
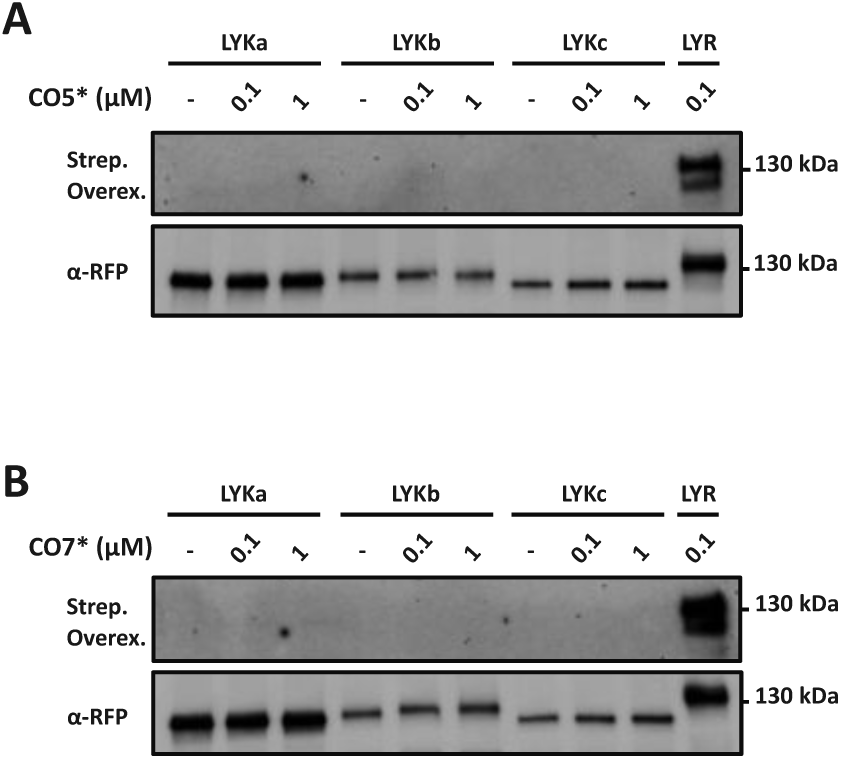
MpaLYR has high affinity for chito- and lipochito-oligosaccharides in *M. paleacea*. **(A)** and (**B**) are identical to the panels shown in **Fig 4B** and **Fig 4D**, respectively, but with over-exposition of the streptavidin-HRP detection. RFP detection was not modified.

**Fig S5.**
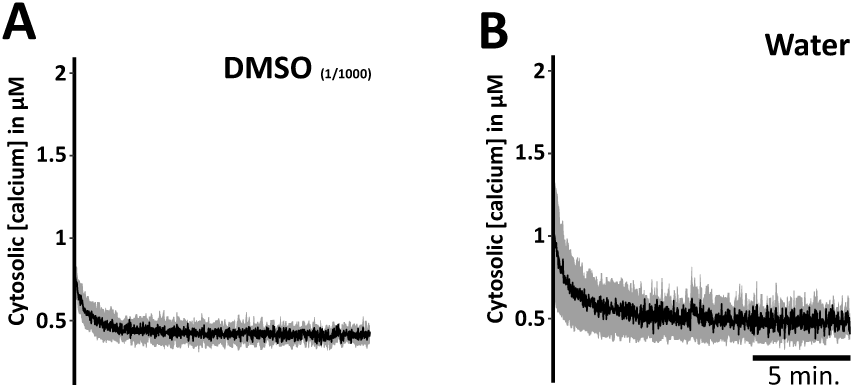
DM5O and water alone do not elicit calcium influx in *M. paleacea*. Cytosolic calcium concentration variations in *M. paleacea* control plants (AEQ-cas9) in response to (**A**) 0.1% dimethyl sulfoxide (DMSO) diluted in water or (**B**) water alone, over a time course of 15 min. (**A-B**) Each trace represents the mean (line) ± standard deviation (shading) from at least three replicates.

**Fig S6.**
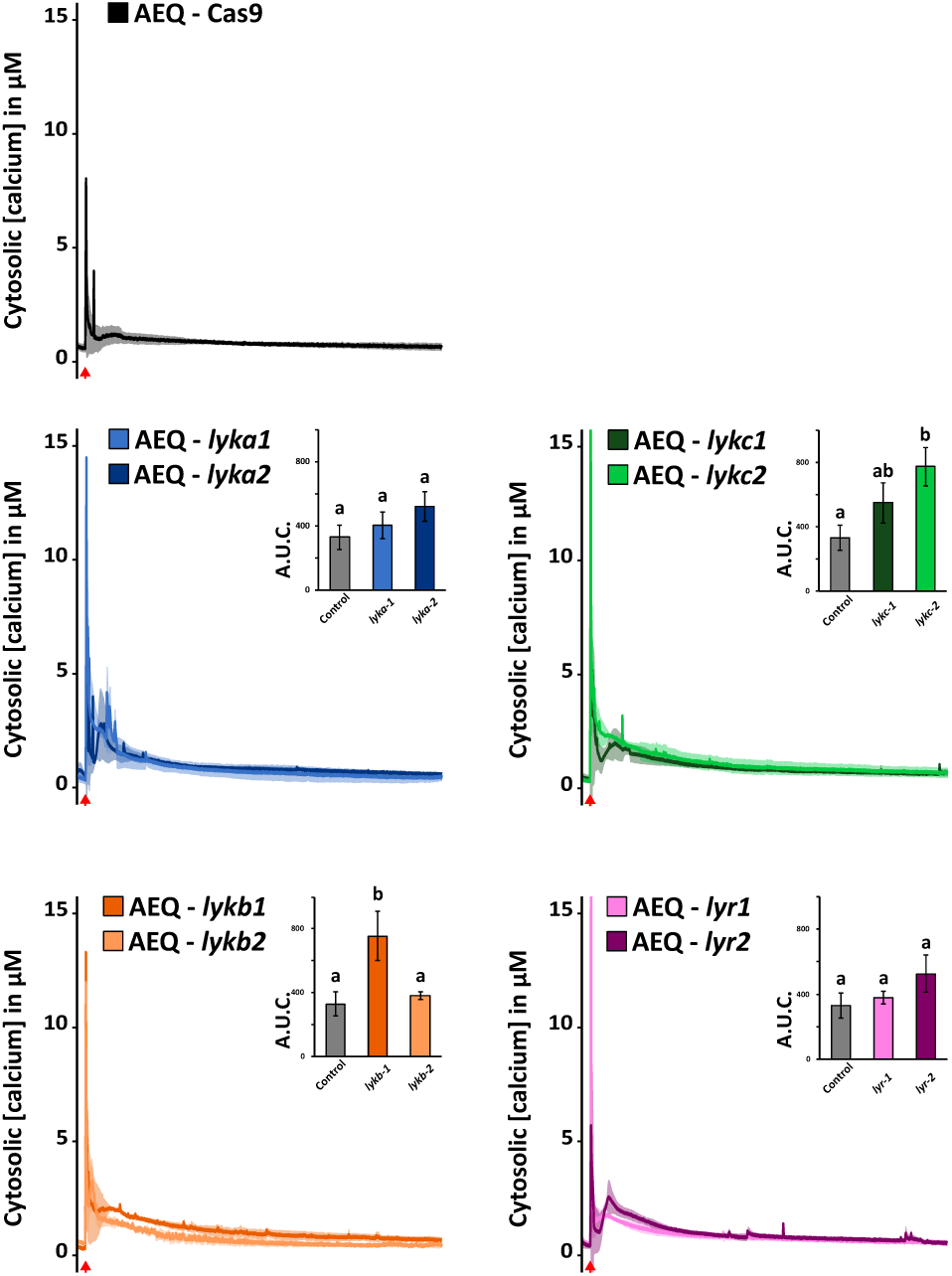
*M. paleacea* LysM-RLK mutants produce a calcium influx in response to hydrogen peroxide. Cytosolic calcium concentration variations in *M. paleacea* control (AEQ-cas9) or two independent mutant lines for each *LysM-RLK*, as indicated, and treated with 1mM hydrogen peroxide diluted in water. Each trace represents the mean (line) ± standard deviation (shading) from three replicates over a time course of 15 min. Bar graph insets represent the area under the curves (A.U.C.) corrected for the baseline and different letters indicate significant differences versus control inferred by ANOVA and Tukey’s HSD post-hoc test (p-value < 0.05).

**Fig S7.**
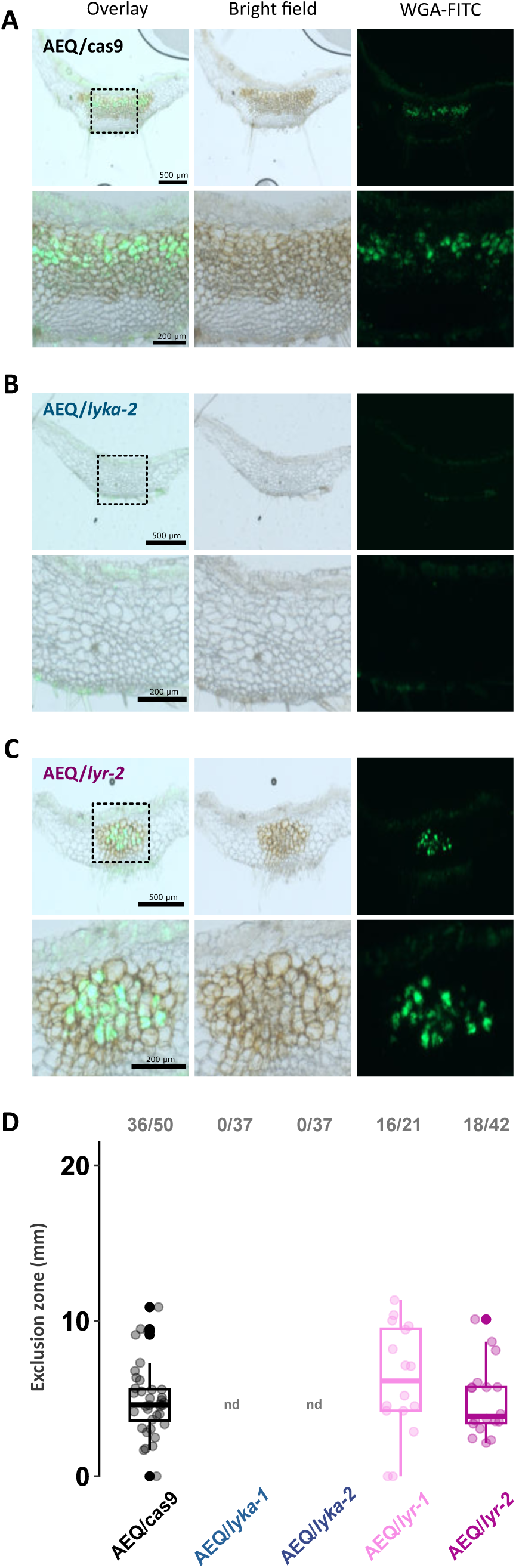
*lyka* and *lyr* mycorrhizal phenotypes in the *M. paleacea* aequorin-expressing background. (**A-C**) Representative images of transversal sections of *M. paleacea* control (AEQ/cas9), one AEQ/*lyka* and one AEQ/*lyr* mutant, six weeks after inoculation with *Rhizophagus irregularis*. Left panels are overlays of bright field (middle panels) and fluorescent (right panels) images. Wheat germ agglutinin (WGA) coupled to Alexa Fluor 488 was used to detect fungal structures. Dashed line delimited insets are enlarged underneath the original images. Scale bars are 500 µm and 200 µm for insets. (**D**) Quantitative analysis of the exclusion zone length on mycorrhizal plants for control (AEQ/cas9) and two independent *lyka* and *lyr* mutants. For each genotype, fractions in bold grey represent mycorrhizal thalli over total thalli assessed.

**Fig S8.**
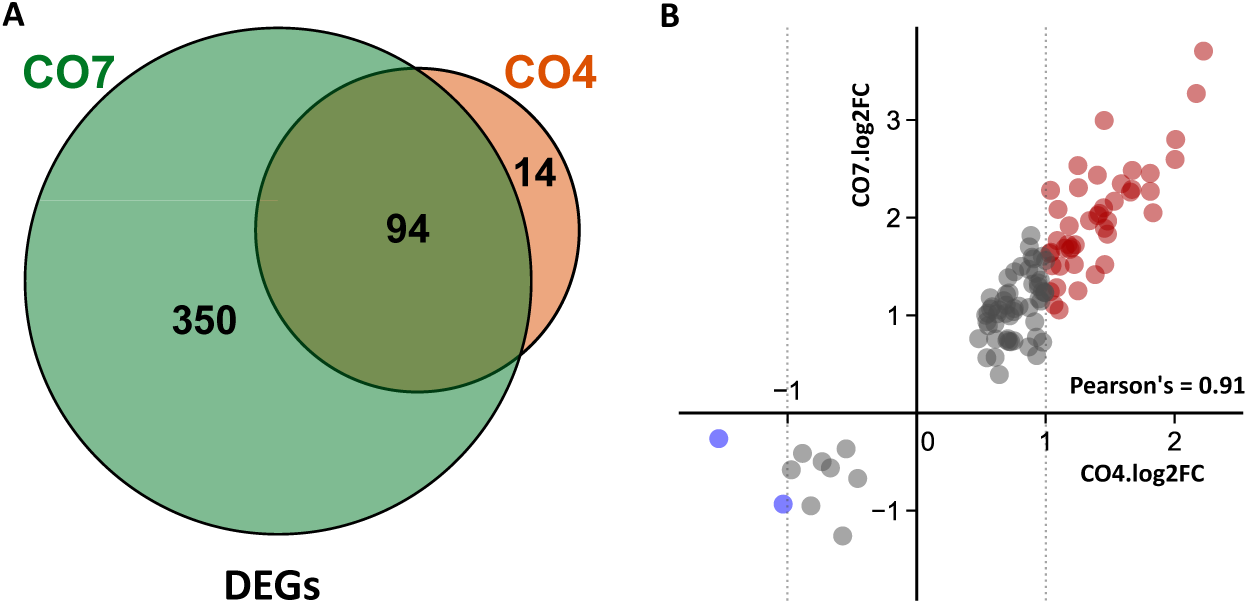
Transcriptomics response to CO4 is a subset of the CO7 response. **(A)** Venn diagram of differentially expressed genes (DEGs) in control plants after one-hour CO7 or CO4 treatment versus water (p adjusted value < 0.05). (**B**) Scatter plot of the log2 fold changes from the 108 DEGs in response to CO4 (x-axis) against their corresponding log2 fold changes in response to CO7 (y-axis). The 45 up-regulated and 2 down-regulated genes in response to CO4 (1log2FC1 ≥ 1 – dashed grey bars) are highlighted in red and blue, respectively. The Pearson’s correlation coefficient between CO7 and CO4 log2FC values is indicated.

**Fig S9.**
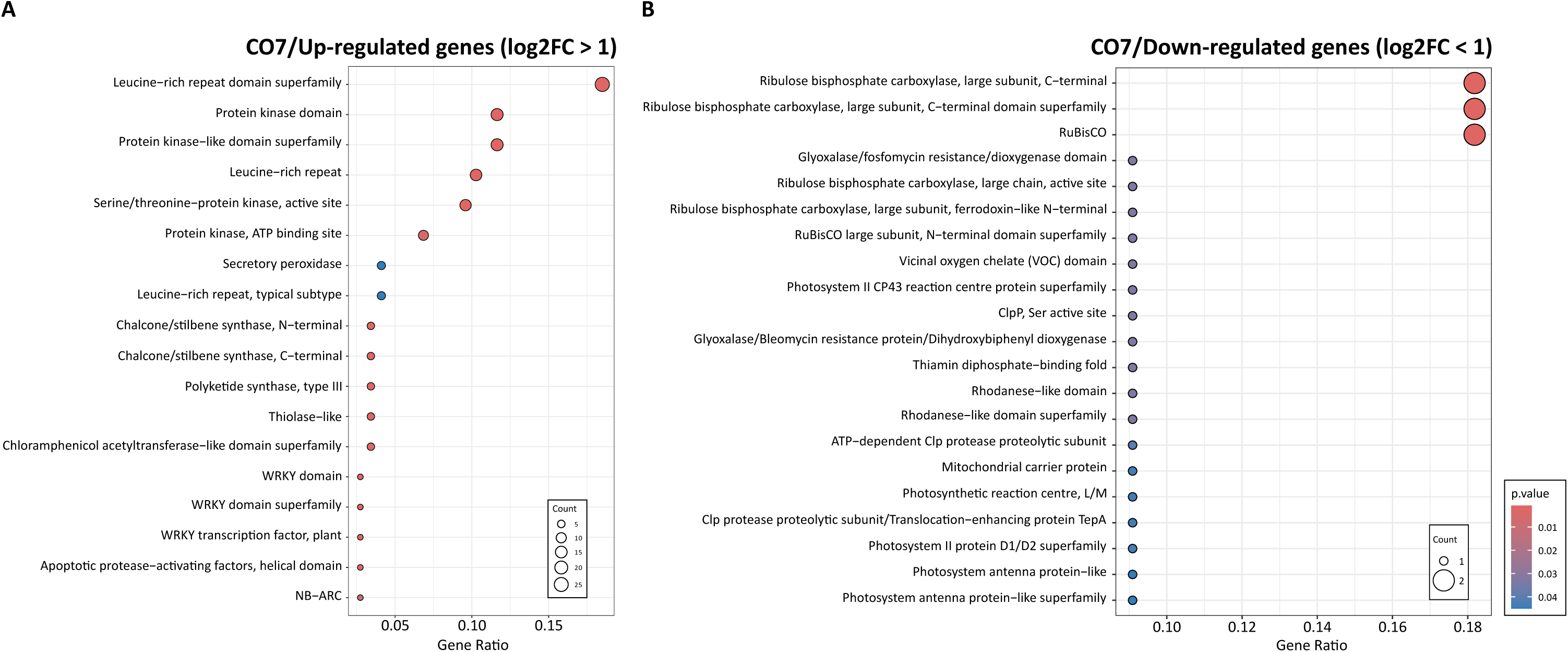
InterPro terms enrichment analyses in response to CO7. InterPro terms enrichment among up-regulated (**A**) and down-regulated (**B**) genes after 1µM CO7 treatment. DEGs are statistically significant (p-value adjusted < 0.05 and 1log2FC1 ≥ 1).

**Fig S10.**
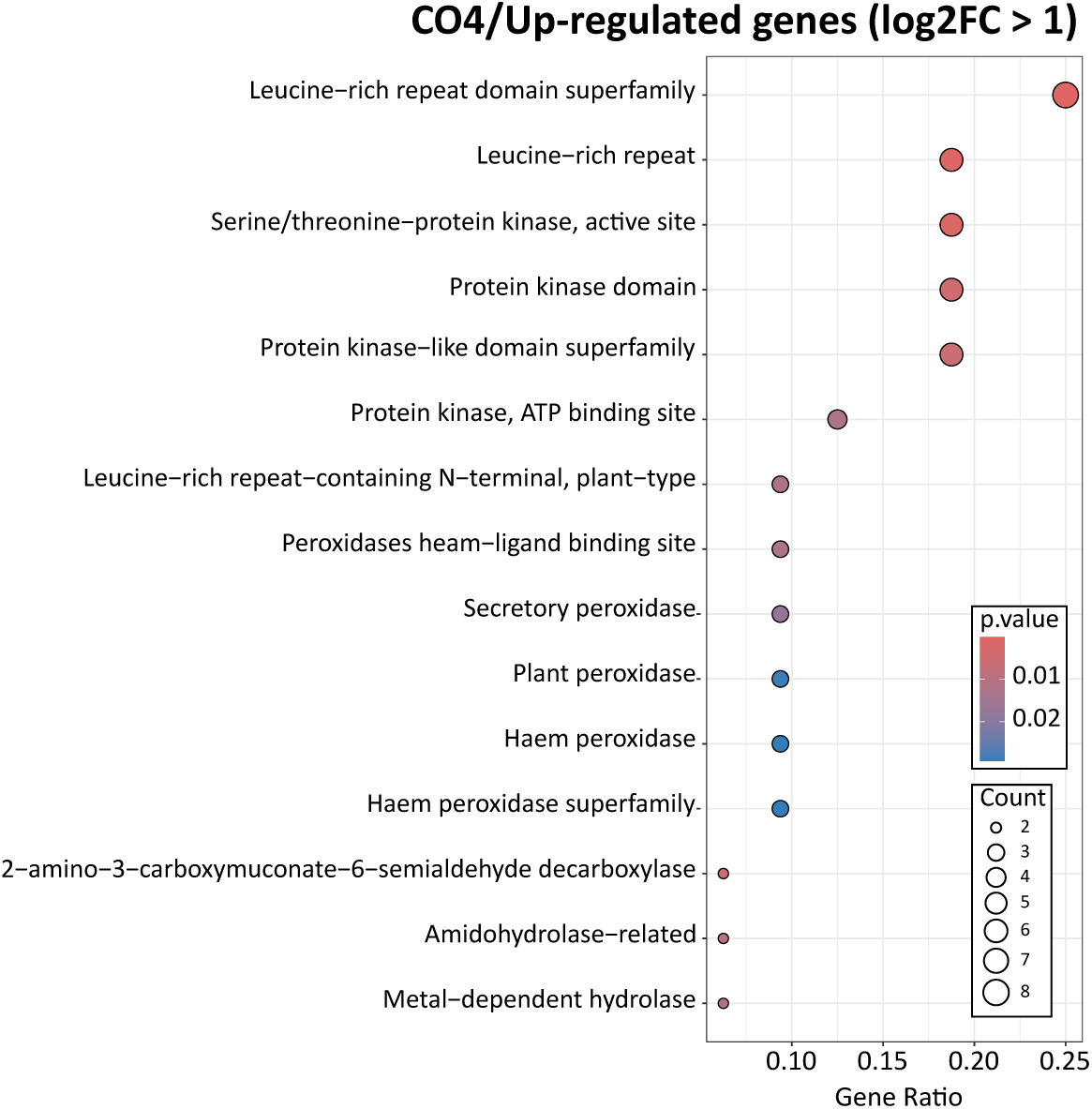
InterPro terms enrichment analysis in response to CO4. InterPro terms enrichment among up-regulated genes after 1µM CO4 treatment. DEGs are statistically significant (p-value adjusted < 0.05 and 1log2FC1 ≥ 1).

**S1 Table.**
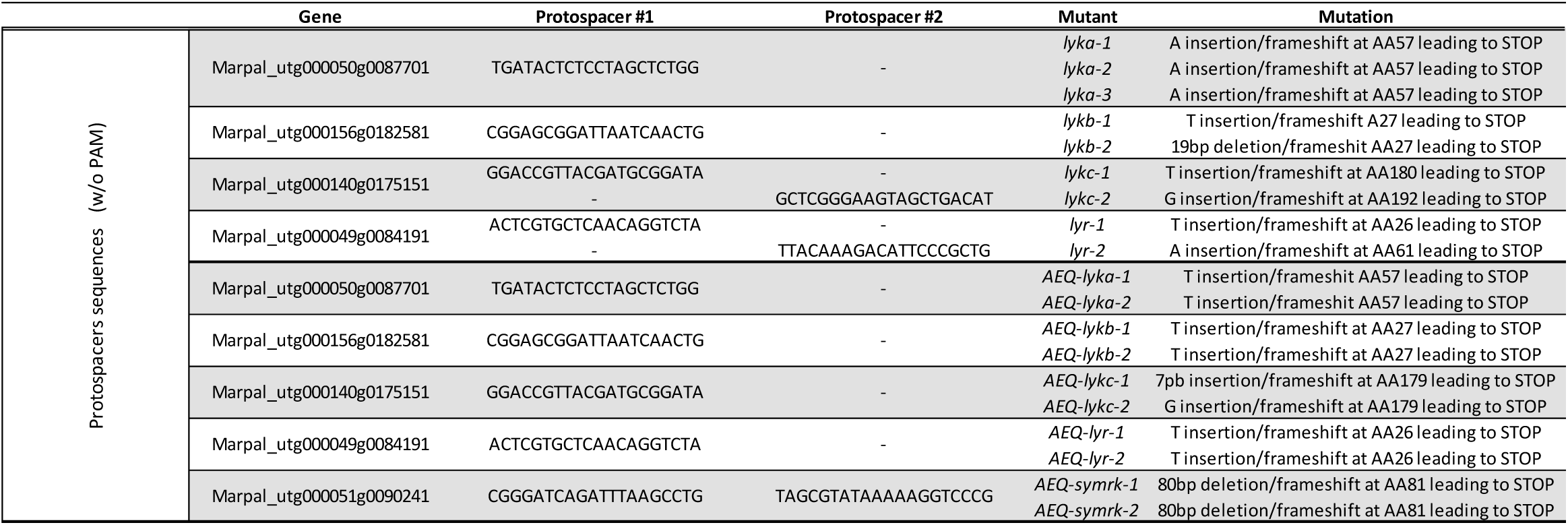
List of protospacers sequences used for gene editions.

S2 Table. List of differentially regulated genes in response to CO7 and CO4 in *M. paleacea* WT and CO7 for *lyka* and *lyr*.

S3 Table. InterPro terms enrichment analyses in response to CO7 and CO4.

**S4 Table.**
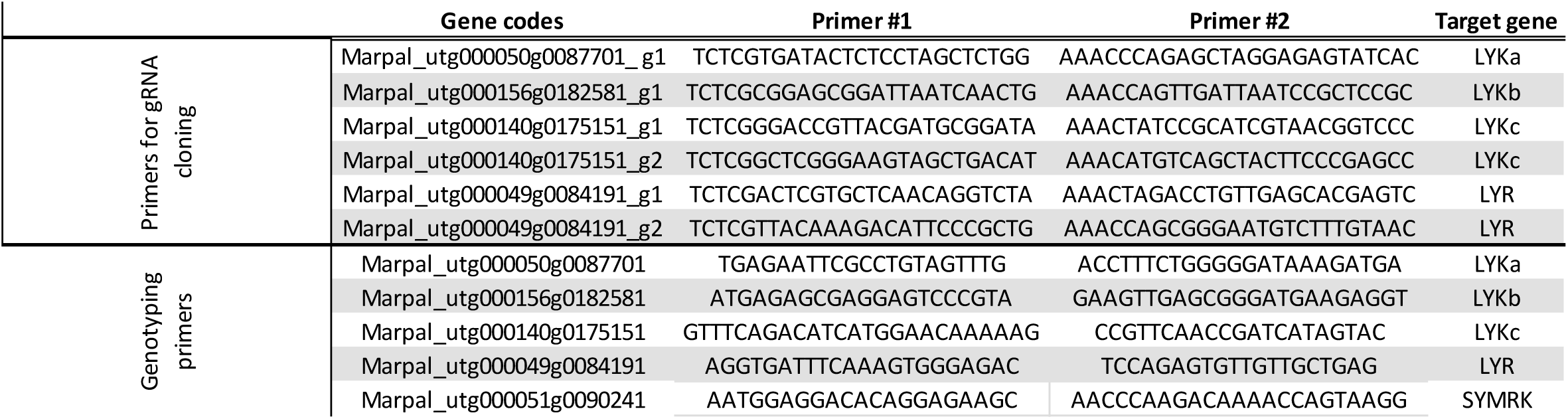
List of primers for genotyping edited lines.

**S5 Table.**
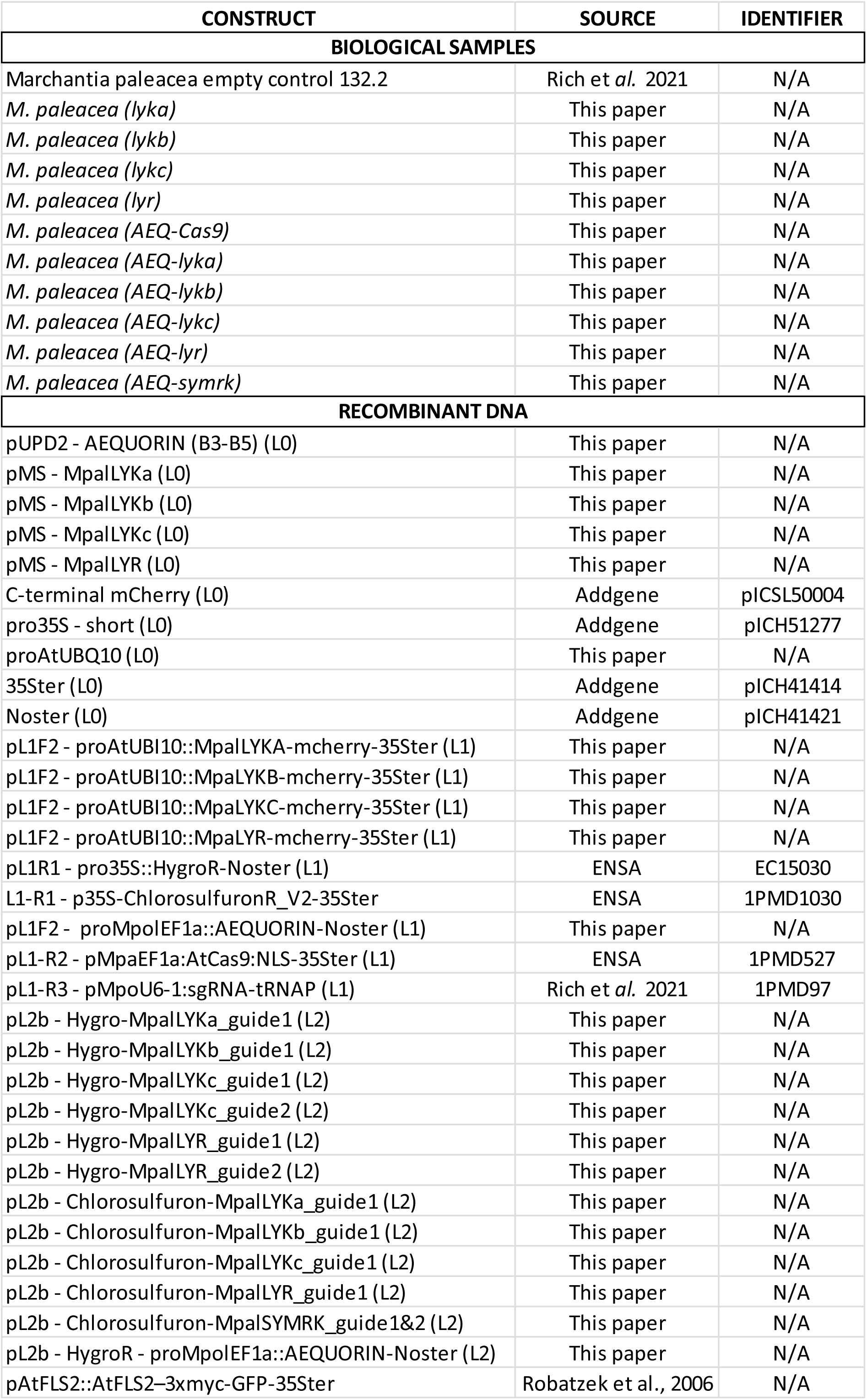
List of constructs and selected lines used in this study.

## Acknowledgements

We would like to thank Vincent Garrigues, Manon Grousset, Lou Battaglia and Madeleine Baker for their technical assistance and Didier Aldon for critically reading the manuscript. E.T. is supported by a PhD grant from the French Ministry of Higher Education and Research. Research performed at LRSV was supported by the ‘Laboratoires d’Excellence (LABEX)’ TULIP (ANR-10-LABX-41) and the ‘École Universitaire de Recherche (EUR)’ TULIP-GS (ANR-18-EURE-0019). P-M.D. is supported by the project Engineering Nitrogen Symbiosis for Africa (ENSA) funded through a grant to the University of Cambridge by the Bill and Melinda Gates Foundation (OPP1172165) and the UK Foreign, Commonwealth and Development Office as Engineering Nitrogen Symbiosis for Africa (OPP1172165) and by funding from the European Research Council (ERC) under the European Union’s Horizon 2020 research and innovation programme (grant agreement no. 101001675 - ORIGINS) to P-M.D. We are grateful to the genotoul bioinformatics platform Toulouse Occitanie (https://bioinfo.genotoul.fr/) for providing computing resources. S.F. acknowledges NanoBio ICMG (UAR 2607) for providing access to mass spectrometry and NMR facilities and the French National Research Agency-ANR through LABEX ARCANE/EUR CBH-EUR-GS (ANR-17-EURE-0003), Glyc@Alps (ANR-15-IDEX-02), and Carnot Polynat (CARN-025-01) for partial financial support.

## Author contributions

E.T. and M.M. designed and coordinated the experiments. S.F. provided material. E.T., S.G., M.R., M.O., D.L., J.K. and M.M. conducted experiments. E.T., D.L, B.L., P.M.D. and M.M. wrote the manuscript. M.M. coordinated the project.

## References

1. Smith SE, Read DJ. Mycorrhizal symbiosis. Elsevier; 2002. doi:10.1016/B978-0-12-652840-4.X5000-1

2. Kodama K, Rich MK, Yoda A, Shimazaki S, Xie X, Akiyama K, et al. An ancestral function of strigolactones as symbiotic rhizosphere signals. Nat Commun. 2022;13: 3974. doi:10.1038/s41467-022-31708-3

3. Kistner C, Parniske M. Evolution of signal transduction in intracellular symbiosis. Trends in Plant Science. 2002;7: 511–518. doi:10.1016/S1360-1385(02)02356-7

4. Stracke S, Kistner C, Yoshida S, Mulder L, Sato S, Kaneko T, et al. A plant receptor-like kinase required for both bacterial and fungal symbiosis. Nature. 2002;417: 959–962. doi:10.1038/nature00841

5. Endre G, Kereszt A, Kevei Z, Mihacea S, Kalo P, Kiss GB. A receptor kinase gene regulating symbiotic nodule development. Nature. 2002;417: 962–966. doi:10.1038/nature00842

6. Miyata K, Hosotani M, Akamatsu A, Takeda N, Jiang W, Sugiyama T, et al. OsSYMRK plays an essential role in AM symbiosis in Rice (*Oryza sativa*). Plant and Cell Physiology. 2023;64: 378–391. doi:10.1093/pcp/pcad006

7. Jin Y, Chen Z, Yang J, Mysore KS, Wen J, Huang J, et al. IPD3 and IPD3L function redundantly in rhizobial and mycorrhizal symbioses. Front Plant Sci. 2018;9: 267. doi:10.3389/fpls.2018.00267

8. Vernie T, Rich M, Pellen T, Teyssier E, Garrigues V, Chauderon L, et al. Conservation of symbiotic signalling across 450 million years of plant evolution. bioRxiv; 2024. p. 2024.01.16.575147. doi:10.1101/2024.01.16.575147

9. Lam AHC, Cooke A, Wright H, Lawson DM, Charpentier M. Evolution of endosymbiosis-mediated nuclear calcium signaling in land plants. Current Biology. 2024;39: 1–9. doi:10.1016/j.cub.2024.03.063

10. Buendia L, Girardin A, Wang T, Cottret L, Lefebvre B. LysM Receptor-Like Kinase and LysM Receptor-Like Protein families: An update on phylogeny and functional characterization. Front Plant Sci. 2018;9: 1531. doi:10.3389/fpls.2018.01531

11. Madsen EB, Madsen LH, Radutoiu S, Olbryt M, Rakwalska M, Szczyglowski K, et al. A receptor kinase gene of the LysM type is involved in legume perception of rhizobial signals. Nature. 2003;425: 637–640. doi:10.1038/nature02045

12. Broghammer A, Krusell L, Blaise M, Sauer J, Sullivan JT, Maolanon N, et al. Legume receptors perceive the rhizobial lipochitin oligosaccharide signal molecules by direct binding. Proc Natl Acad Sci USA. 2012;109: 13859–13864. doi:10.1073/pnas.1205171109

13. Fukuda H, Mamiya R, Akamatsu A, Takeda N. Two L_YS_M receptor-like kinases regulate arbuscular mycorrhiza through distinct signaling pathways in *Lotus japonicus*. New Phytologist. 2024; nph.19863. doi:10.1111/nph.19863

14. Zhang J, Sun J, Chiu CH, Landry D, Li K, Wen J, et al. A receptor required for chitin perception facilitates arbuscular mycorrhizal associations and distinguishes root symbiosis from immunity. Current Biology. 2024; 1–13. doi:10.1016/j.cub.2024.03.015

15. Puttick MN, Morris JL, Williams TA, Cox CJ, Edwards D, Kenrick P, et al. The interrelationships of land plants and the nature of the ancestral embryophyte. Current Biology. 2018;28: 733–745. doi:10.1016/j.cub.2018.01.063

16. Delaux P-M, Hetherington AJ, Coudert Y, Delwiche C, Dunand C, Gould S, et al. Reconstructing trait evolution in plant evo–devo studies. Current Biology. 2019;29: R1110–R1118. doi:10.1016/j.cub.2019.09.044

17. Rich MK, Vigneron N, Libourel C, Keller J, Xue L, Hajheidari M, et al. Lipid exchanges drove the evolution of mutualism during plant terrestrialization. Science. 2021;372: 864–868. doi:10.1126/science.abg0929

18. Delaux P-M, Radhakrishnan GV, Jayaraman D, Cheema J, Malbreil M, Volkening JD, et al. Algal ancestor of land plants was preadapted for symbiosis. Proc Natl Acad Sci USA. 2015;112: 13390–13395. doi:10.1073/pnas.1515426112

19. Nishiyama T, Sakayama H, De Vries J, Buschmann H, Saint-Marcoux D, Ullrich KK, et al. The Chara genome: Secondary complexity and implications for plant terrestrialization. Cell. 2018;174: 448–464. doi:10.1016/j.cell.2018.06.033

20. Yotsui I, Matsui H, Miyauchi S, Iwakawa H, Melkonian K, Schluter T, et al. LysM-mediated signaling in *Marchantia polymorpha* highlights the conservation of pattern-triggered immunity in land plants. Current Biology. 2023;33: 3732–3746. doi:10.1016/j.cub.2023.07.068

21. Miya A, Albert P, Shinya T, Desaki Y, Ichimura K, Shirasu K, et al. CERK1, a LysM receptor kinase, is essential for chitin elicitor signaling in *Arabidopsis*. Proc Natl Acad Sci USA. 2007;104: 19613–19618. doi:10.1073/pnas.0705147104

22. Radutoiu S, Madsen LH, Madsen EB, Felle HH, Umehara Y, Grønlund M, et al. Plant recognition of symbiotic bacteria requires two LysM receptor-like kinases. Nature. 2003;425: 585–592. doi:10.1038/nature02039

23. Radhakrishnan GV, Keller J, Rich MK, Vernie T, Mbadinga Mbadinga DL, Vigneron N, et al. An ancestral signalling pathway is conserved in intracellular symbioses-forming plant lineages. Nat Plants. 2020;6: 280–289. doi:10.1038/s41477-020-0613-7

24. Humphreys CP, Franks PJ, Rees M, Bidartondo MI, Leake JR, Beerling DJ. Mutualistic mycorrhiza-like symbiosis in the most ancient group of land plants. Nat Commun. 2010;1: 103. doi:10.1038/ncomms1105

25. Robatzek S, Chinchilla D, Boller T. Ligand-induced endocytosis of the pattern recognition receptor FLS2 in *Arabidopsis*. Genes Dev. 2006;20: 537–542. doi:10.1101/gad.366506

26. Beck M, Zhou J, Faulkner C, MacLean D, Robatzek S. Spatio-temporal cellular dynamics of the *Arabidopsis* flagellin receptor reveal activation status-dependent endosomal sorting. The Plant Cell. 2012;24: 4205– 4219. doi:10.1105/tpc.112.100263

27. Rush TA, Puech-Pages V, Bascaules A, Jargeat P, Maillet F, Haouy A, et al. Lipo-chitooligosaccharides as regulatory signals of fungal growth and development. Nat Commun. 2020;11: 3897. doi:10.1038/s41467-020-17615-5

28. Zipfel C, Oldroyd GED. Plant signalling in symbiosis and immunity. Nature. 2017;543: 328–336. doi:10.1038/nature22009

29. Knight MR, Read ND, Campbell AK, Trewavas AJ. Imaging calcium dynamics in living plants using semi- synthetic recombinant aequorins. The Journal of Cell Biology. 1993;121.

30. Althoff F, Kopischke S, Zobell O, Ide K, Ishizaki K, Kohchi T, et al. Comparison of the MpEF1α and CaMV35 promoters for application in *Marchantia polymorpha* overexpression studies. Transgenic Res. 2014;23: 235–244. doi:10.1007/s11248-013-9746-z

31. Maillet F, Fournier J, Mendis HC, Tadege M, Wen J, Ratet P, et al. *Sinorhizobium meliloti* succinylated high-molecular-weight succinoglycan and the *Medicago truncatula* LysM receptor-like kinase MtLYK10 participate independently in symbiotic infection. The Plant Journal. 2020;102: 311–326. doi:10.1111/tpj.14625

32. Lewis BD, Spalding EP. Nonselective block by La 3+ of *Arabidopsis* ion channels Involved in signal transduction. Journal of Membrane Biology. 1998;162: 81–90. doi:10.1007/s002329900344

33. Miyata K, Hasegawa S, Nakajima E, Nishizawa Y, Kamiya K, Yokogawa H, et al. OsCERK2/OsRLK10, a homolog of OsCERK1, has a potential role for chitin-triggered immunity and arbuscular mycorrhizal symbiosis in rice. Plant Biotechnology. 2022;39: 119–128. doi:10.5511/plantbiotechnology.21.1222a

34. Miyata K, Kozaki T, Kouzai Y, Ozawa K, Ishii K, Asamizu E, et al. The bifunctional plant receptor, OsCERK1, regulates both chitin-triggered immunity and arbuscular mycorrhizal symbiosis in Rice. Plant and Cell Physiology. 2014;55: 1864–1872. doi:10.1093/pcp/pcu129

35. Gibelin-Viala C, Amblard E, Puech-Pages V, Bonhomme M, Garcia M, Bascaules-Bedin A, et al. The *Medicago truncatula* LysM receptor-like kinase LYK9 plays a dual role in immunity and the arbuscular mycorrhizal symbiosis. New Phytologist. 2019;223: 1516–1529. doi:10.1111/nph.15891

36. Girardin A, Wang T, Ding Y, Keller J, Buendia L, Gaston M, et al. LCO receptors involved in arbuscular mycorrhiza are functional for rhizobia perception in legumes. Current Biology. 2019;29: 4249–4259.e5. doi:10.1016/j.cub.2019.11.038

37. Rutten L, Miyata K, Roswanjaya YP, Huisman R, Bu F, Hartog M, et al. Duplication of symbiotic lysin motif receptors predates the evolution of nitrogen-fixing nodule symbiosis. Plant Physiol. 2020;184: 1004–1023. doi:10.1104/pp.19.01420

38. Gutjahr C, Banba M, Croset V, An K, Miyao A, An G, et al. Arbuscular mycorrhiza–specific signaling in rice transcends the common symbiosis signaling pathway. The Plant Cell. 2008;20: 2989–3005. doi:10.1105/tpc.108.062414

39. Li X-R, Sun J, Albinsky D, Zarrabian D, Hull R, Lee T, et al. Nutrient regulation of lipochitooligosaccharide recognition in plants via NSP1 and NSP2. Nat Commun. 2022;13: 6421. doi:10.1038/s41467-022-33908-3

40. Yano K, Yoshida S, Muller J, Singh S, Banba M, Vickers K, et al. CYCLOPS, a mediator of symbiotic intracellular accommodation. Proc Natl Acad Sci USA. 2008;105: 20540–20545. doi:10.1073/pnas.0806858105

41. Messinese E, Mun J-H, Yeun LH, Jayaraman D, Rouge P, Barre A, et al. A novel nuclear protein interacts with the symbiotic DMI3 calcium- and calmodulin-dependent protein kinase of *Medicago truncatula*. MPMI. 2007;20: 912–921. doi:10.1094/MPMI-20-8-0912

42. Levy J, Bres C, Geurts R, Chalhoub B, Kulikova O, Duc G, et al. A putative Ca ^2+^ and calmodulin-dependent protein kinase required for bacterial and fungal symbioses. Science. 2004;303: 1361–1364. doi:10.1126/science.1093038

43. Catoira R, Galera C, de Billy F, Penmetsa RV, Journet E-P, Maillet F, et al. Four genes of *Medicago truncatula* controlling components of a nod factor ttransduction pathway. The Plant Cell. 2000;12: 1647– 1665.

44. Kawaharada Y, Kelly S, Nielsen MW, Hjuler CT, Gysel K, Muszyński A, et al. Receptor-mediated exopolysaccharide perception controls bacterial infection. Nature. 2015;523: 308–312. doi:10.1038/nature14611

45. Kelly S, Hansen SB, Rubsam H, Saake P, Pedersen EB, Gysel K, et al. A glycan receptor kinase facilitates intracellular accommodation of arbuscular mycorrhiza and symbiotic rhizobia in the legume *Lotus japonicus*. Haney CH, editor. PLoS Biol. 2023;21: e3002127. doi:10.1371/journal.pbio.3002127

46. Roth R, Chiapello M, Montero H, Gehrig P, Grossmann J, O’Holleran K, et al. A rice Serine/Threonine receptor-like kinase regulates arbuscular mycorrhizal symbiosis at the peri-arbuscular membrane. Nat Commun. 2018;9: 4677. doi:10.1038/s41467-018-06865-z

47. Wais RJ, Galera C, Oldroyd G, Catoira R, Penmetsa RV, Cook D, et al. Genetic analysis of calcium spiking responses in nodulation mutants of *Medicago truncatula*. Proc Natl Acad Sci USA. 2000;97: 13407–13412. doi:10.1073/pnas.230439797

48. Kelner A, Leitao N, Chabaud M, Charpentier M, de Carvalho-Niebel F. Dual color sensors for simultaneous analysis of calcium signal dynamics in the nuclear and cytoplasmic compartments of plant cells. Front Plant Sci. 2018;9: 245. doi:10.3389/fpls.2018.00245

49. Shaw SL, Long SR. Nod Factor elicits two separable calcium responses in *Medicago truncatula* root hair cells. Plant Physiology. 2003;131: 976–984. doi:10.1104/pp.005546

50. Binci F, Offer E, Crosino A, Sciascia I, Kleine-Vehn J, Genre A, et al. Spatially and temporally distinct Ca2+ changes in *Lotus japonicus* roots orient fungal-triggered signalling pathways towards symbiosis or immunity. Gifford M, editor. Journal of Experimental Botany. 2024;75: 605–619. doi:10.1093/jxb/erad360

51. Feng F, Sun J, Radhakrishnan GV, Lee T, Bozsóki Z, Fort S, et al. A combination of chitooligosaccharide and lipochitooligosaccharide recognition promotes arbuscular mycorrhizal associations in *Medicago truncatula*. Nat Commun. 2019;10: 5047. doi:10.1038/s41467-019-12999-5

52. Maillet F, Poinsot V, Andre O, Puech-Pages V, Haouy A, Gueunier M, et al. Fungal lipochitooligosaccharide symbiotic signals in arbuscular mycorrhiza. Nature. 2011;469: 58–63. doi:10.1038/nature09622

53. Rasmussen SR, Fuchtbauer W, Novero M, Volpe V, Malkov N, Genre A, et al. Intraradical colonization by arbuscular mycorrhizal fungi triggers induction of a lipochitooligosaccharide receptor. Sci Rep. 2016;6: 29733. doi:10.1038/srep29733

54. Ding Y, Gasciolli V, Medioni L, Gaston M, de-Regibus A, Rem-bliere C, et al. A new group of LysM-RLKs involved in symbiotic signal perception and arbuscular mycorrhiza establishment. biorxiv. 2024 [cited 19 Mar 2024]. doi:10.1101/2024.03.06.583654

55. Miyata K, Hayafune M, Kobae Y, Kaku H, Nishizawa Y, Masuda Y, et al. Evaluation of the role of the LysM Receptor-Like Kinase, OsNFR5/OsRLK2 for AM symbiosis in Rice. Plant Cell Physiol. 2016;57: 2283–2290. doi:10.1093/pcp/pcw144

56. Cullimore J, Fliegmann J, Gasciolli V, Gibelin-Viala C, Carles N, Luu T-B, et al. Evolution of lipochitooligosaccharide binding to a LysM-RLK for nodulation in *Medicago truncatula*. Plant and Cell Physiology. 2023;64: 746–757. doi:10.1093/pcp/pcad033

57. Malkov N, Fliegmann J, Rosenberg C, Gasciolli V, Timmers ACJ, Nurisso A, et al. Molecular basis of lipo-chitooligosaccharide recognition by the lysin motif receptor-like kinase LYR3 in legumes. Biochemical Journal. 2016;473: 1369–1378. doi:10.1042/BCJ20160073

58. Wang T, Gasciolli V, Gaston M, Medioni L, Cumener M, Buendia L, et al. LysM receptor-like kinases involved in immunity perceive lipo-chitooligosaccharides in mycotrophic plants. Plant Physiology. 2023; kiad059. doi:10.1093/plphys/kiad059

59. Cao Y, Liang Y, Tanaka K, Nguyen CT, Jedrzejczak RP, Joachimiak A, et al. The kinase LYK5 is a major chitin receptor in *Arabidopsis* and forms a chitin-induced complex with related kinase CERK1. eLife. 2014;3: e03766. doi:10.7554/eLife.03766

60. Genre A, Chabaud M, Balzergue C, Puech-Pages V, Novero M, Rey T, et al. Short-chain chitin oligomers from arbuscular mycorrhizal fungi trigger nuclear Ca^2+^ spiking in *Medicago truncatula* roots and their production is enhanced by strigolactone. New Phytologist. 2013;198: 190–202. doi:10.1111/nph.12146

61. Genre A, Lanfranco L, Perotto S, Bonfante P. Unique and common traits in mycorrhizal symbioses. Nat Rev Microbiol. 2020;18: 649–660. doi:10.1038/s41579-020-0402-3

62. Cope KR, Bascaules A, Irving TB, Venkateshwaran M, Maeda J, Garcia K, et al. The ectomycorrhizal fungus *Laccaria bicolor* produces lipochitooligosaccharides and uses the common symbiosis pathway to colonize *Populus* roots. Plant Cell. 2019;31: 2386–2410. doi:10.1105/tpc.18.00676

63. Bressendorff S, Azevedo R, Kenchappa CS, Ponce De Le6n I, Olsen JV, Rasmussen MW, et al. An innate immunity pathway in the moss Physcomitrella patens. The Plant Cell. 2016;28: 1328–1342. doi:10.1105/tpc.15.00774

64. Kalyaanamoorthy S, Minh BQ, Wong TKF, von Haeseler A, Jermiin LS. ModelFinder: fast model selection for accurate phylogenetic estimates. Nat Methods. 2017;14: 587–589. doi:10.1038/nmeth.4285

65. Nguyen L-T, Schmidt HA, Von Haeseler A, Minh BQ. IQ-TREE: A fast and effective stochastic algorithm for estimating maximum-likelihood phylogenies. Molecular Biology and Evolution. 2015;32: 268–274. doi:10.1093/molbev/msu300

66. Hoang DT, Chernomor O, Von Haeseler A, Minh BQ, Vinh LS. UFBoot2: Improving the ultrafast bootstrap approximation. Molecular Biology and Evolution. 2018;35: 518–522. doi:10.1093/molbev/msx281

67. Letunic I, Bork P. Interactive Tree Of Life (iTOL) v5: an online tool for phylogenetic tree display and annotation. Nucleic Acids Research. 2021;49: W293–W296. doi:10.1093/nar/gkab301

68. Balzergue C, Puech-Pages V, Becard G, Rochange SF. The regulation of arbuscular mycorrhizal symbiosis by phosphate in pea involves early and systemic signalling events. Journal of Experimental Botany. 2011;62: 1049–1060. doi:10.1093/jxb/erq335

69. Schiessl K, Lilley JLS, Lee T, Tamvakis I, Kohlen W, Bailey PC, et al. NODULE INCEPTION recruits the lateral root developmental program for symbiotic nodule organogenesis in *Medicago truncatula*. Current Biology. 2019;29: 3657–3668.e5. doi:10.1016/j.cub.2019.09.005

70. Lefebvre B, Klaus-Heisen D, Pietraszewska-Bogiel A, Herve C, Camut S, Auriac M-C, et al. Role of N-Glycosylation sites and CXC motifs in trafficking of *Medicago truncatula* Nod Factor perception protein to plasma membrane. J Biol Chem. 2012;287: 10812–10823. doi:10.1074/jbc.M111.281634

71. Li J-F, Norville JE, Aach J, McCormack M, Zhang D, Bush J, et al. Multiplex and homologous recombination– mediated genome editing in *Arabidopsis* and *Nicotiana benthamiana* using guide RNA and Cas9. Nat Biotechnol. 2013;31: 688–691. doi:10.1038/nbt.2654

72. Brini M, Marsault R, Bastianutto C, Alvarez J, Pozzan T, Rizzuto R. Transfected aequorin in the measurement of cytosolic Ca2+ concentration. Journal of Biological Chemistry. 1995;270: 9896–9903. doi:10.1074/jbc.270.17.9896

73. Di Tommaso P, Chatzou M, Floden EW, Barja PP, Palumbo E, Notredame C. Nextflow enables reproducible computational workflows. Nat Biotechnol. 2017;35: 316–319. doi:10.1038/nbt.3820

74. Ewels PA, Peltzer A, Fillinger S, Patel H, Alneberg J, Wilm A, et al. The nf-core framework for community-curated bioinformatics pipelines. Nat Biotechnol. 2020;38: 276–278. doi:10.1038/s41587-020-0439-x

75. Krueger F, James F, Ewels P, Afyounian E, Schuster-Boeckler B. FelixKrueger/TrimGalore: v0.6.7 - DOI via Zenodo. Zenodo; 2021. doi:10.5281/zenodo.5127899

76. Kopylova E, Noe L, Touzet H. SortMeRNA: fast and accurate filtering of ribosomal RNAs in metatranscriptomic data. Bioinformatics. 2012;28: 3211–3217. doi:10.1093/bioinformatics/bts611

77. Quinlan AR, Hall IM. BEDTools: a flexible suite of utilities for comparing genomic features. Bioinformatics. 2010;26: 841–842. doi:10.1093/bioinformatics/btq033

78. Team RC. R: A language and environment for statistical computing. MSOR connections. 2014;1. Available: https://api.semanticscholar.org/CorpusID:215755663

79. Love MI, Huber W, Anders S. Moderated estimation of fold change and dispersion for RNA-seq data with DESeq2. Genome Biology. 2014;15: 550. doi:10.1186/s13059-014-0550-8

80. Sayols S, Scherzinger D, Klein H. dupRadar: a Bioconductor package for the assessment of PCR artifacts in RNA-Seq data. BMC Bioinformatics. 2016;17: 428. doi:10.1186/s12859-016-1276-2

81. Andrews S. FastQC: a quality control tool for high throughput sequence data. 2010 [cited 15 Nov 2024]. Available: http://www.bioinformatics.babraham.ac.uk/projects/fastqc/

82. Pertea G, Pertea M. GFF Utilities: GffRead and GffCompare. F1000Research. 2020;9: ISCB Comm J. doi:10.12688/f1000research.23297.2

83. Wall L, Christiansen T, Orwant J. Programming perl. O’Reilly Media, Inc.; 2000.

84. Van Rossum G, Drake FL. Python 3 Reference Manual. Scotts Valley, CA: CreateSpace; 2009.

85. Li B, Dewey CN. RSEM: accurate transcript quantification from RNA-Seq data with or without a reference genome. BMC Bioinformatics. 2011;12: 323. doi:10.1186/1471-2105-12-323

86. Dobin A, Davis CA, Schlesinger F, Drenkow J, Zaleski C, Jha S, et al. STAR: ultrafast universal RNA-seq aligner. Bioinformatics. 2013;29: 15–21. doi:10.1093/bioinformatics/bts635

87. Picard toolkit. Broad Institute, GitHub repository. Broad Institute; 2019. Available: https://broadinstitute.github.io/picard/

88. Okonechnikov K, Conesa A, Garcia-Alcalde F. Qualimap 2: advanced multi-sample quality control for high-throughput sequencing data. Bioinformatics. 2016;32: 292–294. doi:10.1093/bioinformatics/btv566

89. Wang L, Wang S, Li W. RSeQC: quality control of RNA-seq experiments. Bioinformatics. 2012;28: 2184– 2185. doi:10.1093/bioinformatics/bts356

90. Patro R, Duggal G, Love MI, Irizarry RA, Kingsford C. Salmon provides fast and bias-aware quantification of transcript expression. Nat Methods. 2017;14: 417–419. doi:10.1038/nmeth.4197

91. Morgan M, Obenchain V, Hester J, Pages H. Summarized Experiment: A container (S4 class) for matrix-like assays. Bioconductor. 2024 [cited 15 Nov 2024]. doi:10.18129/B9.bioc.SummarizedExperiment

92. Danecek P, Bonfield JK, Liddle J, Marshall J, Ohan V, Pollard MO, et al. Twelve years of SAMtools and BCFtools. Gigascience. 2021;10: giab008. doi:10.1093/gigascience/giab008

93. Kovaka S, Zimin AV, Pertea GM, Razaghi R, Salzberg SL, Pertea M. Transcriptome assembly from long-read RNA-seq alignments with StringTie2. Genome Biol. 2019;20: 278. doi:10.1186/s13059-019-1910-1

94. Love MI, Soneson C, Hickey PF, Johnson LK, Pierce NT, Shepherd L, et al. Tximeta: Reference sequence checksums for provenance identification in RNA-seq. PLOS Computational Biology. 2020;16: e1007664. doi:10.1371/journal.pcbi.1007664

95. Wu T, Hu E, Xu S, Chen M, Guo P, Dai Z, et al. clusterProfiler 4.0: A universal enrichment tool for interpreting omics data. Innovation (Camb). 2021;2: 100141. doi:10.1016/j.xinn.2021.100141

